# Imaging translational control by Argonaute with single-molecule resolution in live cells

**DOI:** 10.1101/2021.04.30.442135

**Authors:** Charlotte A. Cialek, Tatsuya Morisaki, Ning Zhao, Taiowa A. Montgomery, Timothy J. Stasevich

## Abstract

A major challenge to our understanding of translational control has been deconvolving the individual impact specific regulatory factors have on the complex dynamics of mRNA translation. MicroRNAs (miRNAs), for example, guide Argonaute and associated proteins to target mRNAs, where they direct gene silencing in multiple ways that are not well understood. To better deconvolve these dynamics, we have developed technology to directly visualize and quantify the impact of human Argonaute2 (Ago2) on the translation and subcellular localization of individual reporter mRNAs in living cells. We show that our combined translation and Ago2 tethering system reflects endogenous miRNA-mediated gene silencing. Using the system, we find that Ago2 association leads to progressive silencing of translation at individual mRNA. The timescale of silencing was similar to that of translation, consistent with a role for Ago2 in blocking translation initiation, leading to ribosome runoff. At early time points, we observed occasional brief bursts of translational activity at Ago2-tethered mRNAs undergoing silencing, suggesting that translational repression may initially be reversible. At later time points, Ago2-tethered mRNAs cluster and coalesce with endogenous P-bodies, where a translationally silent state is maintained. These results provide a framework for exploring miRNA-mediated gene regulation in live cells at the single-molecule level. Furthermore, our tethering-based, single-molecule reporter system will likely have wide-ranging application in studying general RNA-protein interactions.

## INTRODUCTION

Translation is the culmination of gene expression, whereby genetic information encoded in nucleic acids is converted into proteins. This basic process is fundamental to all life, giving cells the ability to rapidly establish and maintain diverse phenotypes in response to the environment (Buxbaum et al., 2015; Sonenberg & Hinnebusch, 2009). A multitude of factors work in concert to control which mRNAs are translated, how much peptide product is synthesized, and when to halt erroneous translation (Heck & Wilusz, 2018; Hershey et al., 2019). These regulatory factors are activated by broad cell signaling pathways that respond to stimuli including stress, growth conditions, and development (Roux & Topisirovic, 2018).

The dynamics of translational control have traditionally been studied in living cells at the bulk level (e.g. Western blots, polysome/ribosome profiling) or at the single-cell level (e.g. fluorescent/bioluminescent reporters) (Biswas et al., 2019; Kuersten et al., 2013, 2013). However, these assays lack spatiotemporal resolution, which has made it hard to decipher complex gene regulatory mechanisms (Chao et al., 2012). More recently, it has become possible to explore translation dynamics in living cells at the single-molecule level (Morisaki et al., 2016; Pichon et al., 2016; Wang et al., 2016; Wu et al., 2016; Yan et al., 2016). These single-molecule technologies, which we collectively refer to as Nascent Chain Tracking (NCT), all use repeat mRNA and protein tags to brightly label and track single mRNA using one fluorophore and elongating nascent peptide chains using another fluorophore (reviewed in (Morisaki & Stasevich, 2018)). The ability to image translation at single mRNA makes it possible to quantify individual translation events, measure ribosome initiation and elongation rates, and discern the heterogeneity in translation between mRNA. Since data from this technique are acquired on a microscope, spatial information is naturally embedded. Further, several recent advances in NCT technology have made it possible to study changes in translation at individual mRNA in response to stress (Mateju et al., 2020; Moon, 2020; Moon et al., 2019; Wilbertz et al., 2019), mRNA subcellular location (Voigt et al., 2017), mRNA sequence composition (Aguilera et al., 2019; Boersma et al., 2019; Hoek et al., 2019; Koch et al., 2020; Lyon et al., 2019), and more (as reviewed in (Cialek et al., 2020)).

While powerful, one shortcoming of NCT technology has been visualizing how specific regulatory factors impact translation. Because NCT only amplifies signals from the elongating nascent peptide chain and the mRNA reporter, it is difficult to simultaneously monitor relatively weak signals from individual regulatory factors. Sometimes this difficulty can be avoided if a regulatory factor produces a strong and consistent molecular phenotype that immediately impacts the reporter or its translation. For example, siRNA-directed cleavage of a reporter by Argonaute2 (Ago2) could be detected and quantified using NCT alone because it happened rapidly (seconds to minutes) and resulted in a strong molecular phenotype (the physical splitting of fluorescence signals) (Ruijtenberg et al., 2020). However, translation is often controlled by regulatory factors that act in more subtle ways, for example by inhibiting translation initiation, stalling elongation, or promoting mRNA decay. These modes of regulation tend to have a progressive phenotype that requires extended observation with high sensitivity. To illustrate, in addition to siRNAs, Ago2 binds miRNAs that target mRNAs through partial sequence complementarity and direct gene silencing through a distinct cleavage-independent mechanism (Bartel, 2018). The miRNA-induced silencing complex (miRISC) consists of a miRNA and an Argonaute protein, such as Ago2, along with additional downstream effectors that together promote translational repression and mRNA decay (Jonas & Izaurralde, 2015). Because miRNA-mediated gene silencing and many other gene regulatory mechanisms are likely more gradual and variable than silencing directed by siRNAs, exploring their impact on translation at the single-molecule level is inherently more difficult.

To confront this problem, we developed a “Translation and Tethering” (TnT) single-molecule biosensor that extends NCT technology. As the name implies, the TnT biosensor builds on NCT by adding a tethering cassette that stochastically recruits a fluorescently labeled regulatory factor with controllable stoichiometry. By tracking tethered mRNA through time, the specific and direct impact and regulatory timeframe of the factor can be discerned and quantified. The optimized signal-to-noise ratio of the TnT biosensor makes it possible to continually monitor in 3 colors single-mRNA silencing events that occur on a wide range of timescales, from seconds to hours. Thus, our TnT biosensor provides a route to do highly controlled biochemical experiments with single-molecule spatial resolution and high temporal resolution inside living cells.

Here we introduce the TnT biosensor and demonstrate its functionality by exploring miRNA-mediated gene silencing involving Ago2. We find that within minutes of tethering, Ago2 inhibits the initiation of new translation events, leading to ribosome runoff. Early occurring translational repression is likely independent of mRNA decay, as we sometimes see translation reinitiate in the presence of tethered Ago2. On longer timescales, we find evidence that tethered Ago2 interacts with endogenous miRISC machinery that accumulate at the mRNA and maintain a translationally silent state. Collectively, our data support a model in which Ago2 mediates translational repression largely through reversible inhibition of translation initiation within the cytosol followed by aggregation of mRNA-Ago2 complexes in what are likely P bodies for sustained silencing.

## RESULTS

### A live cell, single-molecule assay to monitor translation and protein tethering in real time

To controllably tether factors to reporter mRNA and simultaneously visualize their impact on mRNA localization, stability, and translation, we constructed a translation and tethering (TnT) biosensor (**Fig. 1A**). As with standard NCT, each TnT biosensor contains 24 MS2 RNA stem loops which recruit fluorescent MS2 Coat Proteins (JF646 HaloTag-MCP) to label the mRNA. Translation is monitored by the localized accumulation of Cy3-conjugated α-FLAG fragmented antibodies (Fab) which can bind 10 FLAG epitopes at the N-terminus of a reporter protein. With this arrangement, as a ribosome begins to translate the TnT biosensor, the nascent protein that emerges from the ribosomal exit tunnel is rapidly labeled by Fab (Morisaki et al., 2016). As multiple ribosomes engage the reporter and translation progresses, the Fab signal intensifies and colocalizes with the mRNA signal. In addition to the mRNA and nascent chain tags, the TnT biosensor also contains a tethering cassette consisting of 15 BoxB RNA stem loops adjacent to the 24 MS2 stem loops in the 3’ UTR. Similar to MCP binding to MS2 stem loops, a 22-amino acid λN protein binds a 19-nucleotide BoxB stem loop with high affinity and specificity (Baron-Benhamou et al., 2004). Thus, by fusing a protein of interest to λN labeled with a distinct fluorophore, it can be recruited and tethered to the BoxB stem loops within an mRNA reporter with high specificity (Coller & Wickens, 2007; Eckhardt et al., 2011). With this arrangement of tags, it is possible to compare individual TnT biosensors that are tethered to a protein of interest to those that are untethered. Furthermore, since the TnT biosensor can theoretically tether up to 15 proteins, the tethering signal is strongly amplified so that tethered mRNA can be tracked for relatively long periods of time without substantial photobleaching or loss of signal within a noisy background.

**Figure 1.**
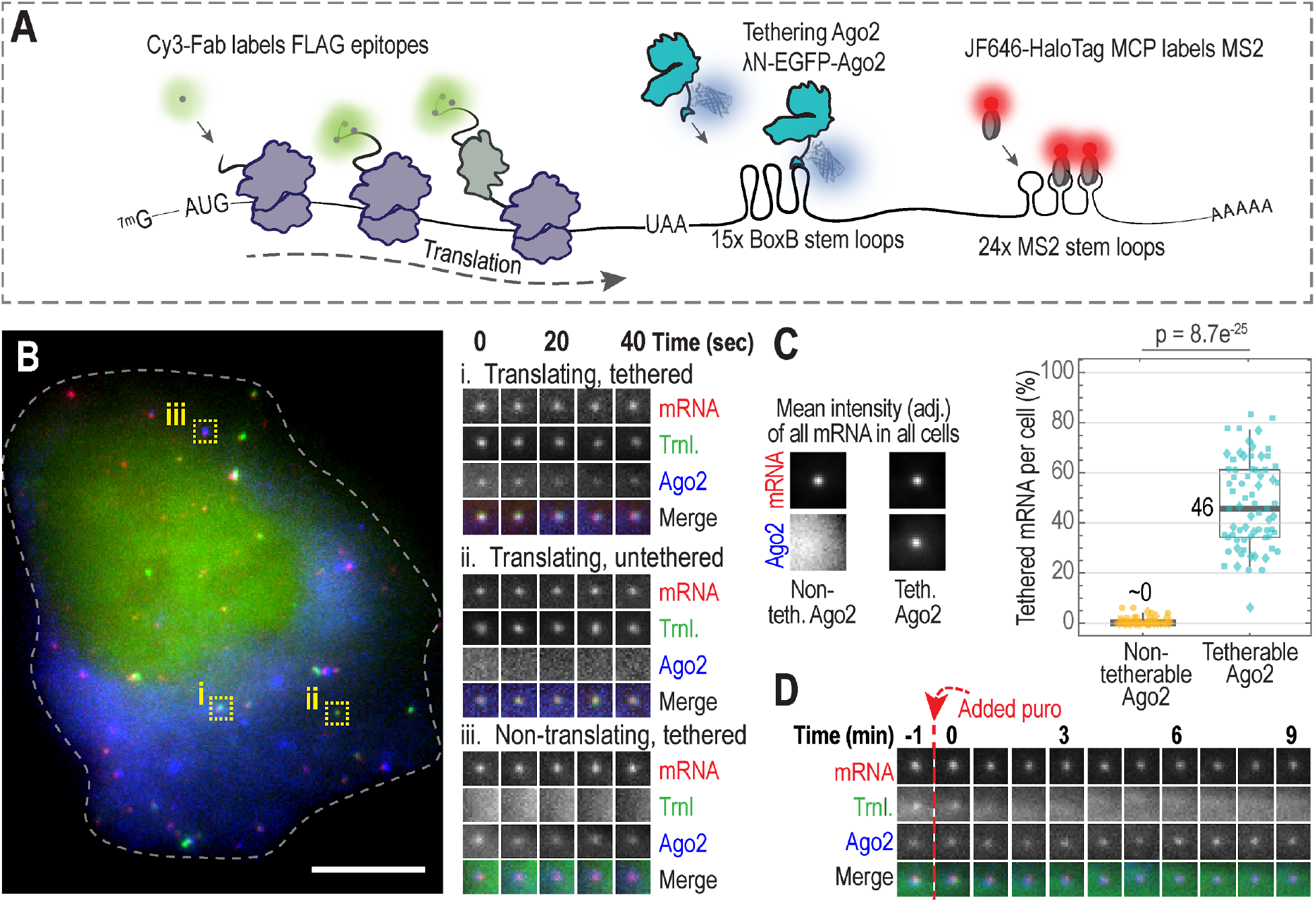
Tracking single-mRNA translation and Argonaute tethering with the TnT biosensor. **A** Schematic of the Translation and Tethering (TnT) biosensor. **B** A representative cell expressing the TnT biosensor to monitor translation (Trnl.) and Ago2-tethering. Single cells were imaged 4 hours after loading plasmids encoding tetherable Ago2 (λN-EGFP-Ago2), the TnT reporter mRNA (smFLAG-KDM5B-15xBoxB-24xMS2), Cy3-FLAG-Fab, and JF646 HaloTag-MCP. The cropped images (15 x 15 pixels^2^; 130 nm/pixel) show single mRNA across a 40 second (sec) interval: (i) translating, tethered; (ii) translating, untethered; and (iii) non-translating, tethered. The dashed line marks the cell outline. Scale bar, 10 μm. **C** The percentages of mRNAs co-localizing with tetherable (λN-EGFP-Ago2) or non-tetherable (EGFP-Ago2) Ago2 6-8 hours after loading was calculated. Left, intensity-rescaled crops showing the average mRNA and Ago2 channels of all detected mRNA foci (18 x 18 pixels^2^; 130 nm/pixel). Right, box plot showing the percentage of mRNA foci colocalizing with Ago2. Each point corresponds to a single cell (shapes denote 3 replicate experiments). The P value was calculated using the Mann-Whitney test. N = 71 (3712) and 65 (7249) cells (mRNA foci) using tetherable and non-tetherable Ago2, respectively. **D** A representative 9 minute (min) track of a single mRNA from a cell (loaded as in B) following puromycin treatment (Time = 0, indicated by the dashed vertical red line). Crops (15 x 15 pixels^2^; 130 nm/pixel) show the mRNA, translation, and Ago2 signal intensities.

To demonstrate the power of the TnT biosensor, we used it to investigate the miRNA-mediated gene silencing pathway. We focused on Ago2, one of the core proteins in the pathway, because others had already shown that tethering it to an mRNA can faithfully recapitulate miRNA-mediated gene silencing (Golden et al., 2017; Liu, Rivas, et al., 2005; Piao et al., 2010; Pillai et al., 2004). While there is a substantial body of evidence that miRNAs regulate gene expression at the level of translation initiation, mRNA decay, and to a lesser extent translation elongation (Reviewed in (Jonas & Izaurralde, 2015)), the timing and spatial organization of these regulatory events and their relative contributions to gene silencing are unclear.

To better deconvolve these complex dynamics, we engineered tetherable (λN-EGFP-Ago2) and non-tetherable (EGFP-Ago2) Ago2 constructs. We then loaded the two Ago2 constructs into human U2OS cells individually along with the other TnT biosensor components (the reporter mRNA, Cy3-labeled α-FLAG Fabs to monitor translation at the reporter, and JF646 HaloTag-MCP to monitor localization of the reporter mRNA). Cells were imaged 4-6 hours after introducing the TnT components. At this time, ~46% of the TnT biosensors colocalized with tetherable Ago2, and some were translationally active (**Fig. 1B,C, Sup. Video 1**). We did not observe non-tetherable Ago2 at the TnT biosensor, indicating that the colocalization we observed was indeed due to tethering (**Fig. 1C and Sup. Fig. 1A**). We next treated cells with the translational inhibitor puromycin. Consistent with the premature release of puromycylated peptide chains during active elongation (Nathans, 1964; Yarmolinsky & Haba, 1959), this led to a rapid loss in translation signals irrespective of whether the cells expressed tetherable or non-tetherable Ago2 (**Fig. 1D, Sup Video 2, and Sup. Fig. 1B,C**). These data therefore demonstrate the functionality of our TnT biosensor, proving it can be used to tether a specific regulatory factor to a trackable mRNA undergoing active translation.

### Ago2 tethering to the TnT reporter mRNA inhibits its translation

Confident in our ability to simultaneously visualize translation and tethering at the single mRNA level, we next explored the impact of Ago2 tethering on translation. First, we set out to confirm that Ago2 tethering reduces total reporter protein synthesis within cells, as previously seen in bulk cell assays (Golden et al., 2017; Liu, Rivas, et al., 2005; Pillai et al., 2004). To control for non-specific silencing that might occur from tethering a protein to the TnT biosensor, we tethered the inert bacterial protein β-gal (λN-EGFP-β-gal), which was previously shown to have a negligible impact on translation (Pillai et al., 2004). Like Ago2, β-gal-tethered mRNAs could be actively translated and were sensitive to inhibition by puromycin (**Sup. Fig. 1D,E**). To control for non-specific effects related to Ago2 overexpression, we also performed experiments with a non-tetherable form of Ago2 (**Fig. 2A**). In each experiment, we quantified total protein production from the TnT biosensor by measuring the accumulation of reporter protein KDM5B (Lysine Demethylase 5B, a nuclear protein) in the nucleus over time (**Fig. 2B**) (Morisaki et al., 2016). According to this metric, cells expressing tetherable Ago2 accumulated ~30% less KDM5B in the nucleus than cells expressing tetherable β-gal or non-tetherable Ago2 (**Fig. 2C**). Importantly, this result was not due to artifactual Fab accumulation in the nucleus (**Sup. Fig. 2A**). As a further test, we repeated this experiment in fixed cells (looking only at a 24 hour time point). Similar to live cells, protein accumulation was reduced 37-55% in fixed cells expressing tetherable Ago2 relative to cells expressing tetherable β-gal or non-tetherable Ago2 (**Sup. Fig. 2B, left**). We included an additional control in this experiment in which a non-tetherable reporter mRNA lacking BoxB stem loops was coexpressed with each tetherable and non-tetherable protein construct. Protein accumulation from this non-tetherable reporter was indistinguishable between the different protein constructs, indicating that Ago2 tethering is directly responsible for the reduction in protein output we observed with the TnT biosensor (**Sup. Fig. 2B, right**). Thus, these data demonstrate that Ago2 tethering leads to a global reduction in mature protein accumulation, consistent with (Golden et al., 2017; Liu, Rivas, et al., 2005; Pillai et al., 2004).

**Figure 2.**
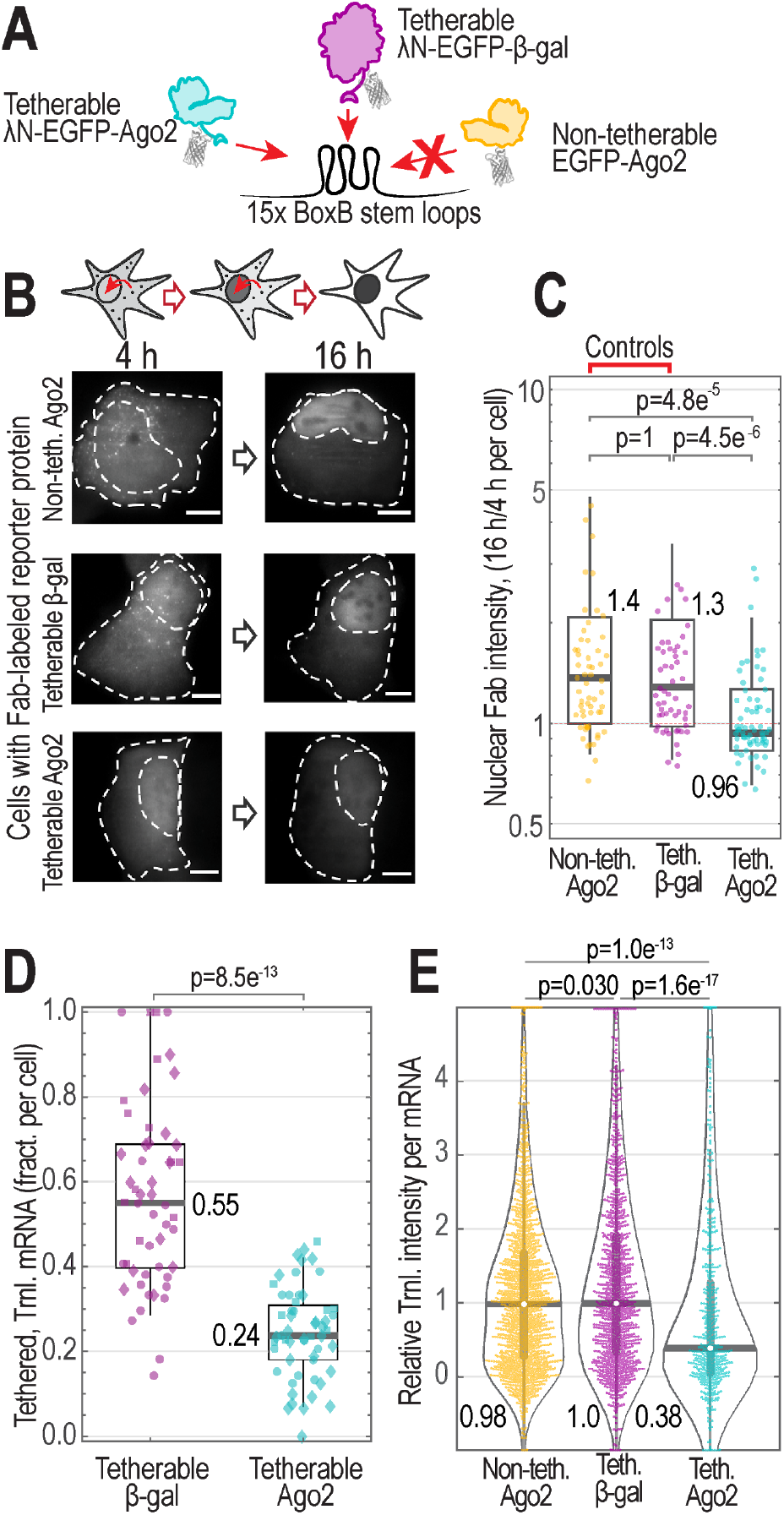
Ago2 tethering represses translation. **A** Schematic of tetherable Ago2 and the controls non-tetherable Ago2 and tetherable β-gal. **B** Schematic (above) and representative cells (below) showing the accumulation of the TnT reporter protein KDM5B (as marked by Fab) in the nucleus over time (4 and 16 hour time points shown) after loading tetherable Ago2 or controls with the TnT components (smFLAG-KDM5B-15xBoxB-24xMS2 mRNA reporter, Cy3-FLAG-Fab, and JF646 HaloTag MCP). Dashed lines mark the cell and nucleus (the nuclear border was defined using cytoplasmic Ago2 or β-gal staining). Scale bars, 10 μm. **C** Box plot displaying data from B. Each data point is the ratio of KDM5B nuclear intensity at 16 hours to 4 hours post-loading of the TnT components. Data from 4 replicate experiments were combined (replicates indicated by marker shape). P values were calculated using the Mann-Whitney test. N = 58 (non-tetherable Ago2), 82 (tetherable β-gal), and 66 (tetherable Ago2) cells. **D** Box plot showing the fraction (frac.) per cell of Ago2- or β-gal-tethered mRNA actively translating 6-8 hours after TnT component loading. Data from 3 replicate experiments were combined (replicates indicated by marker shape). The P value was calculated using the Mann-Whitney test. N = 55 (tetherable β-gal) and 52 (tetherable Ago2) cells. **E** Violin plot displaying TnT reporter translation signal intensity in the presence of tetherable Ago2 or control constructs. The Fab signal intensity was measured for only single mRNA foci. One representative replicate experiment is shown. P values were calculated using the Bonferroni-corrected Mann-Whitney test. N=18 (1513), 28 (1158), 18 (521) cells (mRNA) for non-tetherable Ago2, tetherable β-gal, and tetherable Ago2, respectively.

The main advantage of using the TnT biosensor is the ability to visualize the impact of Ago2 tethering on translation at the individual mRNA level. For this, we reimaged cells expressing both the TnT biosensor and tetherable Ago2 approximately 6 hours after loading, when tethered and translating mRNA could easily be detected (as in **Fig. 1B**). To track individual mRNA for many timepoints, we imaged full cell volumes at a rate of 0.5 frames per second for 20 frames total. Averaging the signals from these short tracks increased the sensitivity and accuracy of translation and tethering detection. To control for tethering and Ago2 overexpression artifacts, we again imaged control cells loaded with either non-tetherable Ago2 or tetherable β-gal. For each condition, we quantified the number of mRNA that were tethered/untethered and translating/silent (**Sup. Fig. 2C**).

In line with the low levels of mature reporter protein accumulation we observed in the nucleus, cells expressing tetherable Ago2 had, on average, the fewest mRNAs being translated (**Sup. Fig. 2C**). In fact, just ~24% of Ago2-tethered mRNAs were being translated compared to ~55% of β-gal-tethered mRNAs (**Fig. 2D**). There was also a 30-50% reduction in the total number of mRNA foci per cell using tetherable Ago2 (**Sup. Fig. 2C**), indicating a shortened mRNA half-life, although we could not rule out the reduction was in part caused by mRNA clustering and transcriptional stochasticity. Finally, individual mRNA in cells expressing tetherable Ago2 had translation signals that were 40-60% as bright as those in cells expressing tetherable β-gal (**Fig. 2E and Sup. Fig. 2D**). Taken together, these data demonstrate that Ago2 tethering not only reduces the fraction of mRNA being translated, but also reduces the number of translating ribosomes on individual mRNAs.

### Ago2-tethered mRNA coalesce in the cytoplasm

Due to the exceptional signal-to-noise of the TnT biosensor, we could monitor individual mRNA in living cells for extended periods of time, ranging from seconds to hours. By following mRNA in single cells over a period of 12+ hours, we noticed that translationally silenced, Ago2-tethered mRNA tended to cluster over time (**Fig. 3A**). These mRNA clusters remained translationally silenced and colocalized with bright cytoplasmic Ago2 foci for hours. Notably, they also exhibited behaviors characteristic of phase-separation, such as coalescence (**Sup. Video 3**). The average intensity of mRNA foci in cells expressing tetherable Ago2 steadily increased with time (**Fig. 3B**). In contrast, the average intensity of mRNA foci in cells expressing tetherable β-gal or non-tetherable Ago2 slightly decreased with time, likely due to photobleaching or probe decay (**Fig. 3B**).

**Figure 3.**
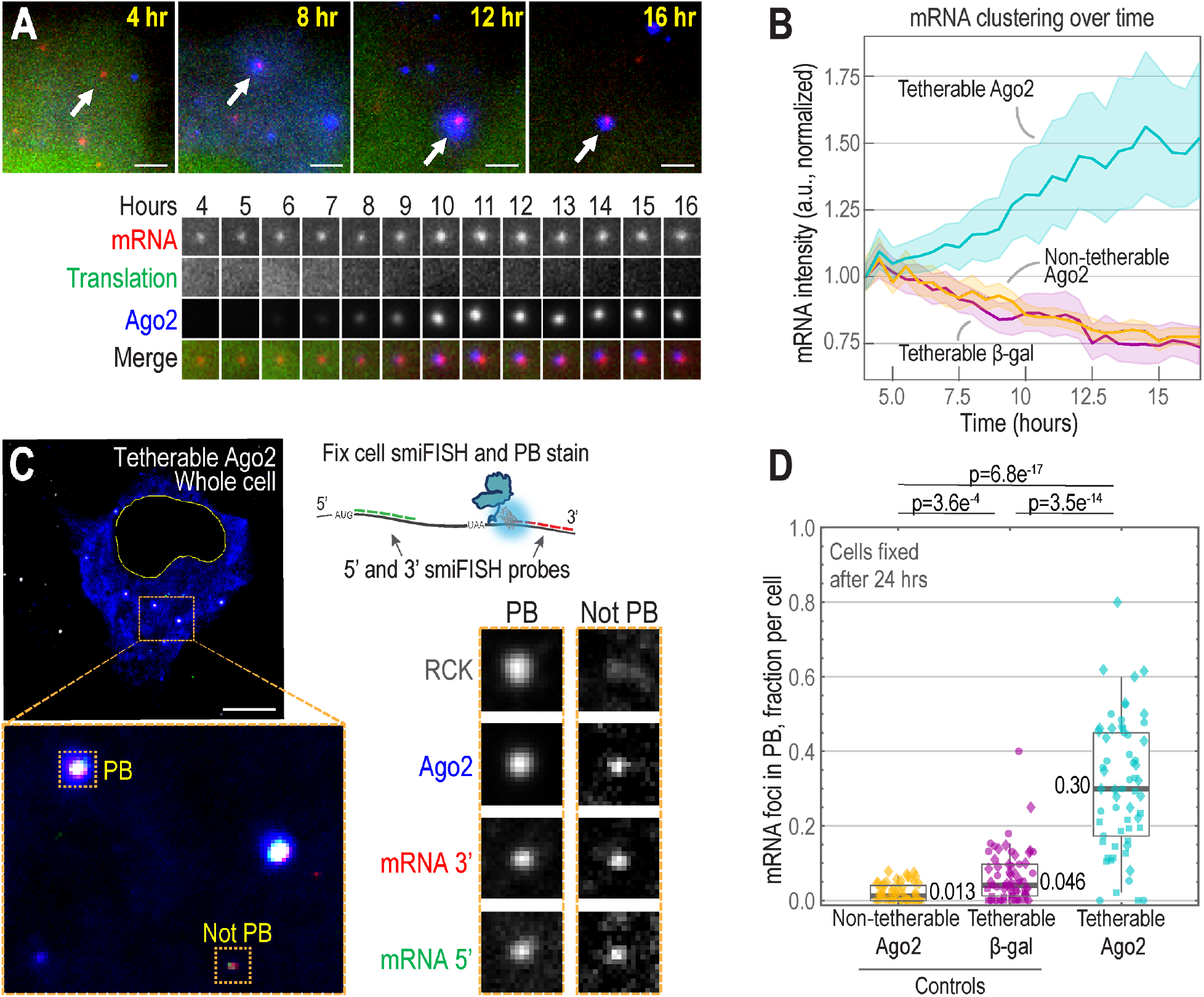
Intact Ago2-tethered TnT biosensors cluster and accumulate in P-bodies after translational silencing. **A** A representative cytoplasmic patch from a live cell expressing tetherable Ago2 and the TnT biosensor and components (smFLAG-KDM5B-15xBoxB-24xMS2 mRNA reporter, Cy3-FLAG-Fab, and JF646 HaloTag MCP) was imaged 4-16 h post loading. The mRNA identified by arrows is shown over time below in crops (15 x 15 pixels^2^; 130 nm/pixel). Scale bars, 2.5 μm. **B** The line plot displays mRNA clustering over time. Cells were imaged 4-16 h post loading with the TnT biosensor plus either tetherable Ago2, tetherable β-gal, or non-tetherable Ago2. The mean MCP signal of all mRNA in all cells was measured over time and normalized to 1 at the first timepoint. N = 35 (17,367), 46 (32,666) and 44 (19,236) cells (mRNA foci) for non-tetherable Ago2, tetherable β-gal, and tetherable Ago2, respectively. Shaded region, 95% CI. **C-D** The TnT biosensor plasmid plus either the tetherable Ago2, tetherable β-gal, or non-tetherable Ago2 plasmid were loaded into cells. After 24h, cells were fixed. The 5’ and 3’ mRNA ends were labeled with smiFISH probes, and P-bodies were stained with α-RCK antibodies. **C** Representative tetherable-Ago2 cell, cytoplasmic patch, and mRNAs displaying P-body (PB) localization. Colors show P-bodies (α-RCK, gray; not shown on left), expressed tetherable Ago2 (blue), and TnT mRNA smiFISH probes (5’, green; 3’, red). Crops (17 x 17 pixels^2^; 130 nm/pixel) were autoscaled. Scale bar, 10 μm. **D** Box plots showing the fraction per cell of mRNA with 5’ and 3’ smFISH signals that were colocalized with P-bodies. Each point represents the fraction from a single cell (yellow, non-tetherable Ago2; purple, tetherable β-gal; cyan, tetherable Ago2; different shapes mark 1 of 3 replicate experiments). N = 56 (3,655), 63 (3,958), 61 (2,999) cells (mRNA foci) for non-tetherable Ago2, tetherable β-gal, and tetherable Ago2, respectively. P values were calculated from Bonferroni-corrected Mann-Whitney tests.

Based on these observations, we wondered if Ago2 tethering caused the TnT biosensor to relocalize to specific subcellular locations or compartments. Ago2 is known to interact directly with TNRC6 (Elkayam et al., 2017; Liu, Rivas, et al., 2005; Sheu-Gruttadauria & MacRae, 2018) and indirectly with other downstream effectors in the miRISC complex, including factors required for P-body assembly, such as RCK/DDX6 (Ayache et al., 2015; Chen et al., 2014; Rouya et al., 2014; Standart & Weil, 2018). Thus, we hypothesized progressive mRNA clustering could arise from multivalent interactions with endogenous miRISC machinery and/or P-bodies. Consistent with our observations of coalescing mRNAs, this machinery can exhibit phase-separation behavior both *in vitro* and *in vivo* (Lin et al., 2015; Moon et al., 2019; Pitchiaya et al., 2019; Sheu-Gruttadauria & MacRae, 2018).

To test this hypothesis, we costained fixed cells expressing the TnT biosensor with smiFISH probes (Tsanov et al., 2016) complementary to the 3’ and 5’ ends of mRNA in separate colors and with an α-RCK antibody to label P-bodies (although we note RCK foci could also be in complex with Ago2 outside of P-bodies). As we hypothesized, the bright mRNA-Ago2 foci did indeed colocalize with RCK (**Fig. 3C**). Further, the localization was dependent on Ago2-tethering. About 30% of mRNA colocalized with RCK in cells expressing tetherable Ago2, compared to less than 5% in control cells (**Fig. 3D**).

Interestingly, the majority of mRNAs we observed contained both 5’ and 3’ smiFISH signals, regardless of tethering or colocalization with RCK (**Sup. Fig. 3A**). While this would suggest that at least some mRNAs in P-bodies are intact, as seen previously (Horvathova et al., 2017), it was difficult to know for sure since each cluster contained many mRNAs and the 5’ and 3’ signals could therefore come from different molecules. Furthermore, since the distinct smiFISH probes bind with different affinities and have fluorophores with distinct properties, it was also difficult to say if the 5’ and 3’ signals were present at a one-to-one stochiometry, although we did observe a similar ratio of signals at mRNA both inside and outside of RCK foci (**Sup. Fig. 3B**).

Collectively, these data show that tethering Ago2 to an mRNA is sufficient to recruit endogenous downstream factors involved in miRNA-mediated translational repression. Furthermore, the recruitment we observed can lead to unique mRNA behaviors, including long-term translational silencing, as well as mRNA clustering and coalescence reminiscent of phase-separation. More generally, these data demonstrate how the TnT biosensor can be used to target mRNA to specific subcellular locations or microenvironments to better understand how they affect translation dynamics.

### Progressive loss of translation upon Ago2 tethering

Thus far we have shown that Ago2-tethered TnT biosensors are translationally silenced and ultimately cluster into RCK foci that are presumably P-bodies. With this in mind, we next sought to zoom in on the kinetics of translation silencing by tracking single TnT biosensors prior to their clustering. According to several lines of work, Ago2 is thought to silence mRNA in part by inhibiting ribosome initiation (Jonas & Izaurralde, 2015)). However, direct visual evidence for this has been lacking due to the limited spatiotemporal resolution of earlier experiments. Assuming ribosome initiation is inhibited, we would expect the TnT translation signal to dim after an Ago2 tethering event. Dimming would be gradual as elongating ribosomes finish translating the open reading frame and run off the transcript one by one. Other mechanisms of translational repression are also possible and can be discerned using the TnT biosensor. For example, if ribosomes prematurely abort translation after Ago2 tethering, the translation signal would rapidly disappear, as occurs when cells are treated with puromycin (**Fig. 1D, Sup. Fig. 1B,C,E**). Also, if translation elongation is repressed, ribosome progression could be slowed or halted, in which case the translation signal would persist beyond the time it takes to translate the open reading frame. Finally, if an mRNA were sliced, as in siRNA-mediated translational silencing, this would result in the physical separation of the translation and mRNA signals (Ruijtenberg et al., 2020).

To distinguish between these possibilities, we developed an imaging strategy to track freely diffusing TnT biosensors with high temporal resolution for upwards of 90 minutes. Specifically, we imaged whole cell volumes every 10 seconds in the mRNA channel, allowing us to track a single mRNA for well over an hour. Concurrently, we imaged translation and tethering once every 100 seconds. This staggered approach allowed us to follow the translation and tethering signals of individual mRNAs with minimal photobleaching. From 11 cells over 5 days, we identified 25 mRNA that were trackable for 35-90 minutes and that had at some point both tethering and translation signals. We repeated the experiment in 11 cells expressing tetherable β-gal over 4 days, finding 21 such control-tethered mRNA.

For each tracked mRNA, we measured the intensities of the mRNA, translation, and tethering signals through time. Consistent with an inhibition of translation initiation and ribosome runoff, we could observe a slow and steady decline of the translation signal from individual Ago2-tethered mRNA (**Fig. 4A, Sup. Video 4**). Moreover, a scatterplot of translation signals versus tethering signals for all data points revealed a striking pattern: translation signals were strongest when Ago2 tethering signals were weakest, and vice versa (**Fig. 4B, cyan**). In stark contrast, this inverse relationship between translation and tethering signals (Spearman correlation coefficient = −0.10; p = 1.7×10^−4^) was not observed when mRNA were tethered to control β-gal (**Fig. 4B, purple**). In fact, β-gal tethering signals were correlated with translation (Spearman correlation coefficient = 0.55; p = 2.4×10^−81^). This suggested that the more Ago2 present, the less likely it is that the mRNA is actively translating. To visualize this over time, we normalized all tracked signals and averaged them together (so each mRNA would have equal weight) (**Fig. 4C**). This revealed translation signals steadily and significantly decreased with time (t_1/2_ ~40 minutes), while Ago2 tethering signals steadily increased (**Fig. 4C, right**). The corresponding signals in control cells expressing tetherable β-gal remained steady (**Fig. 4C, left**). These data strongly suggest Ago2 tethering leads to a gradual runoff of ribosomes, as would be expected if ribosomal initiation were inhibited.

**Figure 4.**
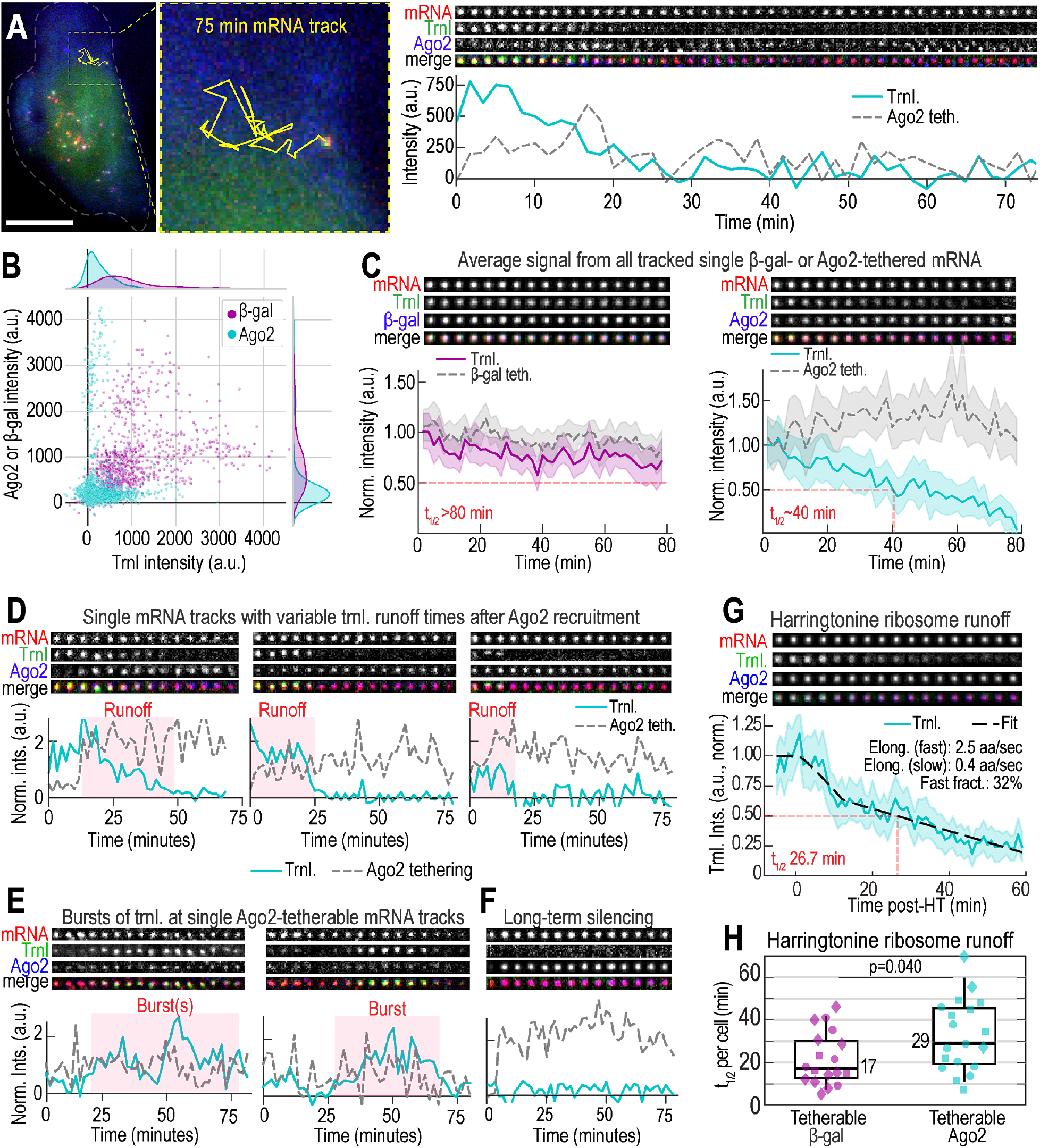
Progressive loss of translation upon Ago2 tethering. **A** A representative track of a single TnT biosensor in a cell expressing tetherable Ago2 and loaded with TnT components (smFLAG-KDM5B-15xBoxB-24xMS2 mRNA reporter, Cy3-FLAG-Fab, and JF646 HaloTag MCP) over a 75-minute interval. Left, the track of the mRNA (yellow line) is overlaid on the cell image. Right, crops (11 x 11 pixels^2^; 130 nm/pixel) of this track are displayed over time to show the mRNA, translation (Trnl), and Ago2 tethering signals every 100 seconds. These signals are plotted over time below (Trnl, solid cyan line; Ago2, dashed gray line). Scale bar, 10 μm. **B** Scatter plot showing single-mRNA tethering signals (β-gal, purple; Ago2, cyan) versus translation signals (Trnl) for all tracked TnT biosensors at all timepoints (β-gal, 21 tracks, 11 cells; Ago2, 25 tracks, 11 cells). The Spearman Correlation Coefficient was calculated from this data (0.55 for β-gal, p=2×10^−81^; −0.10 for Ago2, p=2×10^−4^). **C** Line plots, as in A, showing average signals from multiple TnT biosensor tracks through time. Signals from individual β-gal or Ago2 tetherable tracks were normalized from 0 to 1 and then averaged, so no one track would dominate. The resulting average curves were then renormalized (using the first four timepoints) so they started from ~1. Left, average translation (Trnl.) and tethering (Teth.) signals from cells expressing tetherable β-gal (β-gal-teth., dashed gray line; Trnl, solid purple line). Right, average translation and tethering signals from cells expressing tetherable Ago2 (Ago2-teth., dashed gray line; Trnl, solid cyan line). Shaded regions, 95% CIs. Solid red lines denote the runoff halftime t_1/2_. **D** Sample tracks of single TnT biosensors (renormalized as in B; every 3rd crop is shown after a 3-frame rolling average). Runoff times shorten from left to right. **E** Sample tracks of single TnT biosensors showing bursts of translation, plotted as in D. **F** Sample track showing long-term silencing with high Ago2 tethering signals, plotted as in D. **G** Average single-molecule signals from all detected TnT biosensors in a single representative cell expressing tetherable Ago2 and exposed to HT (Time = 0), plotted as in C. The average translation signal (solid line) showing ribosome runoff was fit to a biphasic model (dashed line; fitted parameters displayed). Red line denotes the runoff halftime t_1/2_. **H** Box plot showing all fitted single-cell HT-runoff halftimes. N=18 cells expressing tetherable Ago2; N = 17 cells expressing tetherable β-gal (marker shapes denote 3 replicate experiments).

We next examined the individual mRNA tracks in greater detail. In line with the average behavior in **Fig. 4C**, the translation signals at many steadily declined as in **Fig. 4A**, although the rate of runoff was variable from mRNA to mRNA (**Fig. 4D**). This variability most likely reflected stochasticity in the start of our imaging as well as in the positioning of individual ribosomes along an mRNA upon Ago2 tethering. In particular, we noticed runoffs could be delayed by late Ago2 tethering (**Fig. 4D, left**) and runoffs with low initial translation signal intensities could be faster (**Fig. 4D, middle vs. right**), presumably because they were already in progress when imaging began. We also observed occasional bursts of translation despite the presence of low-levels of Ago2 tethering (**Fig. 4E**). This would suggest the repression of translation initiation by Ago2 is reversible. Further, we observed Ago2-tethered, translationally-silent mRNA (**Fig. 4F, Sup. Video 5**) coalesce to form large clusters that were similar to the P-bodies we identified in **Fig. 3**. These mRNA were associated with exceptionally bright Ago2 foci that likely contained many Ago2 proteins. Finally, on two occasions we observed physical separation of the translation and mRNA signals, possibly indicating some form of mRNA cleavage (Horvathova et al., 2017; Ruijtenberg et al., 2020) (**Sup. Fig. 4A, Sup. Video 6**). Given the scarcity of this type of event (2/25; <8% of observations), we conclude it is not a dominant form of translation repression in our system.

### Translational repression at Ago2-tethered mRNA is consistent with inhibition of translation initiation

Our analysis of the individual mRNA tracks above provides further support for a model in which Ago2 tethering strongly inhibits translation initiation. To further confirm this model, we performed Harringtonine experiments (HT). HT inhibits translation initiation while allowing already-initiated, elongating ribosomes to continue translation and run off the transcript (Huang, 1975). If Ago2 tethering inhibits initiation as potently as HT, we would predict Ago2-tethered ribosome runoff times should be on the same timescale as HT-induced runoff times.

To test this prediction, we added HT to cells expressing the TnT biosensor and either tetherable Ago2 or tetherable β-gal. This led to a steady loss in the average translation signal from mRNA. In cells expressing tetherable Ago2, the HT-induced ribosome runoff halftime was ~29 minutes, with the majority of signal lost in ~60 minutes (**Fig. 4G and Sup. Video 7**). This timescale is similar but a bit faster than what we observed when tracking individual Ago2-tethered mRNA in the absence of HT (**Fig. 4C, right**). These data indicate Ago2-tethering is nearly as strong at inhibiting translation initiation as HT.

Interestingly, in cells expressing tetherable β-gal, the HT-induced runoff appeared faster (**Fig. 4H and Sup. Fig. 4B**). This overall faster runoff time could be seen in the average, photobleach-corrected runoff curves from multiple experiments, although the difference was not always as significant due to high cell-to-cell variability (**Sup. Fig. 4C,D**). To quantify the difference, we fit each single-cell HT runoff curve to a biphasic model (dashed line in **Fig. 4G and Sup. Fig. 4B**) in which a fraction of elongating ribosomes run off with relatively fast kinetics and the remaining fraction run off with relatively slow kinetics (see **Methods** for model details). According to the individual fits, the Ago2 slowdown arises because there are fewer ribosomes in the faster elongating state (~29% vs. ~45%; **Sup. Fig. 4E-G**). These results therefore provide evidence that Ago2 not only inhibits translation initiation, but also leads to a moderate but measurable slowdown in elongation.

### Ago2 tethering mimics miRNA-mediated translational repression at the single-molecule level

Although previous work provided convincing evidence that Ago2 tethering recapitulates natural miRNA-mediated translational silencing (Eckhardt et al., 2011; Golden et al., 2017; Liu, Rivas, et al., 2005; Pillai et al., 2004), we wanted to confirm this at the single-molecule level to further validate our TnT biosensor data. To achieve this, we created modified TnT biosensors containing endogenous miRNA response elements (MREs) in place of the tethering cassette. For MREs, we chose the 3’UTR of the POLR3G gene, which is predicted by TargetScan (*TargetScanHuman 7.2*, n.d.) to contain three endogenous MREs targeted by miR-26-5p. To confirm these MREs repress translation in our cells, we first placed them in the 3’UTR of a simpler reporter that encodes sfGFP-H2B (**Sup. Fig. 5A**). Cells loaded with the MRE-containing reporter produced ~40% less sfGFP-H2B after 24 hours than cells loaded with a reporter containing mutated MREs, indicating that the intact MREs promote miRNA-mediated silencing (**Sup. Fig. 5B**).

**Figure 5.**
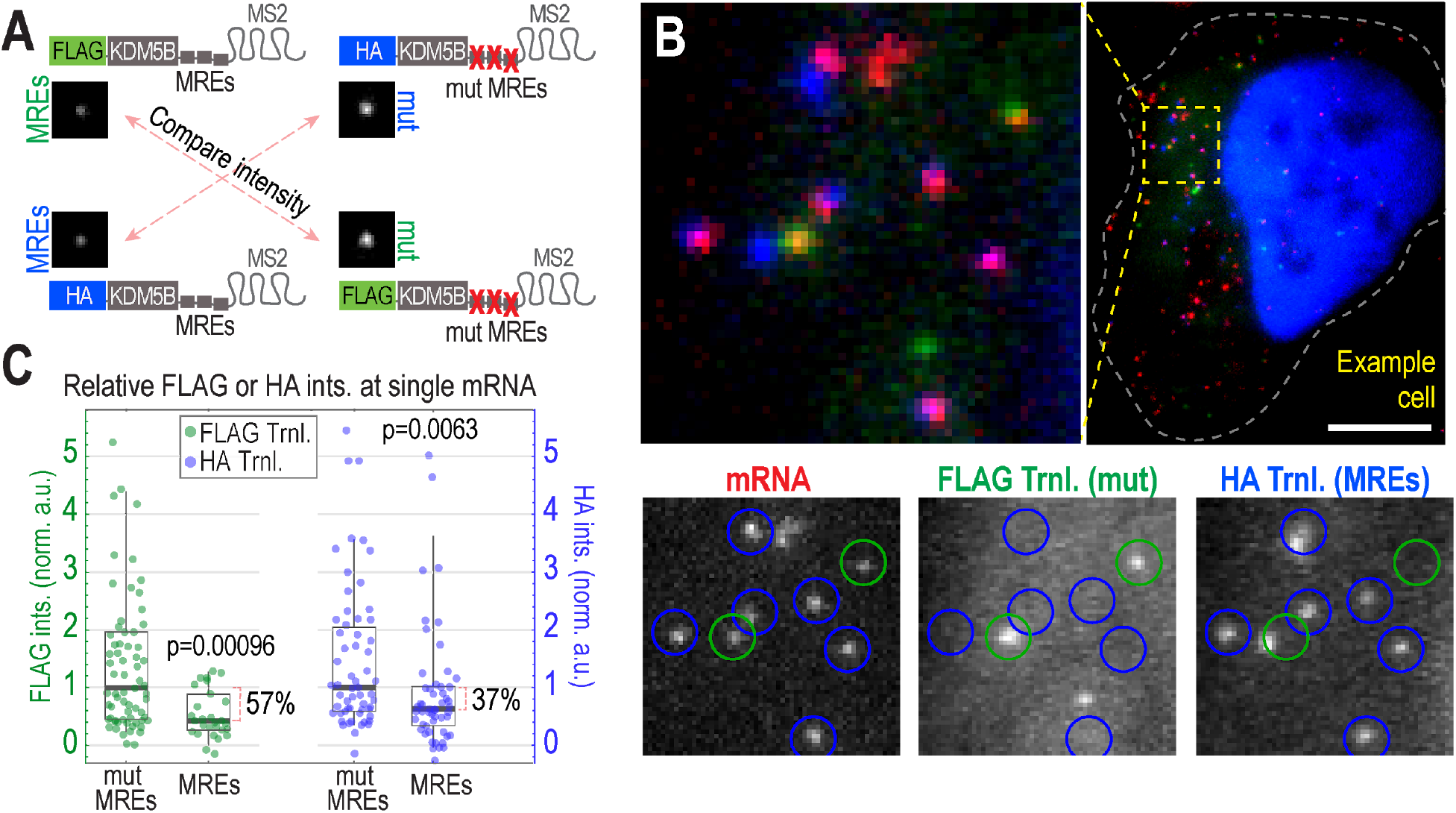
miRNA-directed Ago targeting represses translation at the single-mRNA level. **A** Schematic of modified TnT biosensors with either endogenous miRNA Response Elements (MREs) or inactive, mutated versions (MUT) inserted in place of the tethering cassette. In each experiment, two modified biosensors (either top two or bottom two) were coexpressed in single cells. To fairly gauge relative translation signals, FLAG (or HA) signals were compared across experiments (red, dashed arrows). **B** An example cell loaded with the top two biosensors in A. Image above shows a whole cell and a representative cytoplasmic patch. Image below shows each separated channel from the patch (intensity auto-adjust ed), with detected translating (Trnl) mRNA circled in blue (indicating translation of HA) or green (indicating translation of FLAG). Scale bar, 10 μm. **C** Box plots showing the relative intensity (ints.) of the mutated MREs (mut MREs) and endogenous MREs (MREs) containing modified biosensors from 1 of 3 replicate experiments (see **Sup. Fig. 5C** for the other two replicate experiments). Each point represents the measured translation signal from a single detected biosensor (N=69 mut MREs and N=30 MREs for FLAG; N=58 mut MREs and N=54 MREs for HA). P values were calculated using the Mann-Whitney test.

We then introduced the MRE cassette in place of the BoxB hairpins in our TnT biosensor as well as a similar biosensor that encoded 10 HA instead of 10 FLAG epitopes. In addition, we introduced the mutated MREs into both of these biosensors (**Fig. 5A**). These miRNA-based biosensors provide a more natural context than the TnT biosensor, but with the drawback that we can no longer track Ago2 association. Nevertheless, the miRNA-based biosensors can still be tracked at the single-molecule level, allowing us to measure translation at individual reporter foci. Assuming our original TnT biosensor recapitulates natural miRNA-mediated silencing, we would predict that our new biosensors containing endogenous MREs would on average have lower translation signals (presumably due to ribosome runoff) than those containing mutated MREs.

To test this, we coloaded cells with one of two combinations of biosensors: either a FLAG miRNA-biosensor with endogenous MREs and an HA miRNA-biosensor with mutated MREs, or the reciprocal pair of biosensors in which FLAG and HA were swapped (**Fig. 5A**). This allowed us to directly compare endogenous and mutant MRE-containing biosensors in the same cells (**Fig. 5B, Sup. Video 8**). We then quantified the translation signals at individual biosensors in live cells. Translation was reduced by 38-57% (across 3 replicates) in cells containing our FLAG-based biosensors with endogenous MREs relative to cells containing the mutant versions and by 0-54% (across 3 replicate) in the reciprocal situation with HA-based biosensors (**Fig. 5C, Sup. Fig. 5C**). Thus, in agreement with our prediction, endogenous MREs do significantly reduce the translation signals of individual reporter mRNAs. Moreover, the overall 37-57% reduction in translation in all but one replicate is in good agreement with the 40-60% reduction we observed when tethering Ago2 to the original TnT biosensor (**Fig. 2E**). Taken together, these data provide additional single-molecule evidence that tethering Ago2 to the TnT biosensor recapitulates endogenous miRNA-mediated Ago2 translational repression.

## DISCUSSION

The Translation and Tethering (TnT) biosensor reported here allows for direct visualization of specific regulatory factors and quantification of their impact on translation at individual mRNA molecules in living cells. The TnT biosensor innovation builds off of Nascent Chain Tracking technology, adding an independent tethering cassette composed of repeat BoxB stem loops in the 3’ UTR of the biosensor. This makes it possible to track individual mRNA molecules and simultaneously monitor their translational status both before and after tethering events. Since tethering is highly specific and the signal amplified, the impact of regulatory factors on translation can be quantified and their actions on timescales ranging from seconds to hours deconvolved. The TnT biosensor therefore brings us one step closer to performing controlled, single mRNA biochemical reactions in the natural environment of living cells.

To demonstrate the applicability of the TnT biosensor, we used it to deconvolve the complex translational regulatory dynamics of Ago2. There has been a long-term debate about the precise temporal ordering of translational silencing by Ago2 (Jonas & Izaurralde, 2015). Multiple models have been proposed, some stating sequential translational repression and mRNA decay, others stating decay occurs co-translationally, and still others stating silencing and decay can be uncoupled (Bazzini et al., 2012; Béthune et al., 2012; Bose et al., 2017; Djuranovic et al., 2012; Eichhorn et al., 2014; Guo et al., 2010; Pillai et al., 2005; Subtelny et al., 2014; Tat et al., 2016; Wilczynska et al., 2019). According to our data, within minutes of Ago2 tethering to an mRNA, translation initiation is strongly inhibited, leading to a gradual loss of translating ribosomes as they run off the transcript. Collectively, our data support a model in which the inhibition of translation initiation is an early step in the miRNA-mediated silencing pathway, one that can act independently of other steps, such as mRNA decay. Two lines of evidence support this model: first, translational bursts could occur after an mRNA was silenced by Ago2 tethering (**Fig. 4E**); second, it appeared that nearly all TnT biosensors had 5’ and 3’ ends according to smiFISH (**Sup. Fig. 3**). Together, this evidence suggests an mRNA that has been translationally silenced by Ago2 tethering can remain intact and later be translated. Thus, inhibition of translation initiation can be decoupled from mRNA decay, at least in our tethering system.

In addition to the clear impact Ago2 tethering had on translation initiation, we also measured a moderate impact on elongation (**Fig. 4G,H and Sup. Fig. 4B,D,G**). While it remains unclear how this slight slowdown in elongation would impact translational silencing, in principle the slowly moving ribosomes would increase the probability of incidental ribosomal collisions (Hickey et al., 2020; Juszkiewicz et al., 2020). If this were to happen, these collisions can be sensed by quality control machinery. According to recent work from the Hegde and Green labs, this machinery can inhibit translation initiation so that additional ribosomal collisions do not occur and elongation can proceed (Hickey et al., 2020; Juszkiewicz et al., 2020). While it remains unclear if this sort of coupling between the repression of elongation and initiation is occurring in our system, it could be one more contributing factor in Ago2-dependent translational silencing.

By tracking the TnT biosensor over longer timescales, we also discovered both Ago2 and mRNA signals progressively increased with time. The buildup in Ago2 signals began immediately, during ribosome runoff (**Fig. 4C, right**), while the buildup in mRNA signals occurred over many hours as mRNA and Ago2 molecules coalesced (**Fig. 3A,B**). The nature of this progressive accumulation of signals is consistent with diverse, multivalent interactions with endogenous miRISC machinery. In particular, a single tethered Ago2 protein can theoretically bind up to three TNRC6B proteins via three distinct binding pockets (Schirle & MacRae, 2012; Sheu-Gruttadauria & MacRae, 2018). Likewise, a single TNRC6A protein can bind up to three Ago2 proteins via three distinct binding sites (Elkayam et al., 2017). In turn, TNRC6 proteins contain long, unstructured domains that can recruit other, more downstream miRISC effectors, such as deadenylases, decapping complexes, and translational repression factors like RCK/DDX6 (Jonas & Izaurralde, 2015). Such diverse, multivalent protein-protein and protein-RNA interactions are hallmarks of phase separation (Lin et al., 2015; Moon et al., 2019; Pitchiaya et al., 2019; Sheu-Gruttadauria & MacRae, 2018), and minimal Ago2-TNRC6 complexes have in fact been observed to phase separate both *in vivo* and *in vitro* (Lin et al., 2015; Meister et al., 2005; Pillai et al., 2005; Sen & Blau, 2005), leading to increased sequestration of miRNA target mRNAs and deadenylases (Sheu-Gruttadauria & MacRae, 2018). It therefore seems likely the TnT biosensor serves as a proxy for these large RNA-protein complexes. While the presence of RCK would suggest some are mature P-bodies, it is possible that Ago2 and RCK interact outside of P-bodies. The slow and progressive accumulation in mRNA and Ago2 we observed would imply that our TnT biosensor offers a live-cell, single-molecule window into the seeding of phase-separated bodies.

Our experimental design was based off of earlier work from the Filipowicz and Parker labs, which pioneered the use of the Ago2 tethering system to investigate miRNA-directed Ago2 translational silencing (Eckhardt et al., 2011; Liu, Valencia-Sanchez, et al., 2005; Pillai et al., 2004, 2005). Using the TnT biosensor, we corroborated their finding that tethering Ago2 to a reporter mRNA leads to a global reduction in reporter protein synthesis (**Fig. 2B,C and Sup. Fig. 2B**). Furthermore, by comparing the TnT biosensor to a more natural one with endogenous MREs, we demonstrated that Ago2 tethering remains a good model of miRNA-directed translational silencing, even at the single-molecule level (**Fig. 5 and Sup. Fig. 5**).

This work is similar to a companion study by Kobayashi and Singer (will cite) that developed a different miRNA-based reporter to investigate translational silencing by Ago2 with single-molecule precision *in situ*. Their measurements also indicate Ago2 silences mRNA translation on the 30-40 minute timescale, prior to mRNA decay. Although the two reporter systems are similar, a unique advantage of ours is the amplified tethering signal, which allowed us to track the dynamics of Ago2-tethered mRNA for long periods of time in living cells.

Despite these reassuring results, there are three important caveats of our approach that we should point out. First, the artificial tethering of Ago2 to mRNA bypasses the natural interaction mediated by miRNA base-pairing. This could misorient tethered Ago2 so that it does not behave in a completely natural way. Nonetheless, the consistency of translation silencing between the TnT biosensor and the more natural miRNA-based biosensors we tested suggests that tethered Ago2 retains its core silencing functionality. Second, our tethering cassette was quite large, requiring 15 BoxB stem loops to track tethering for extended periods of time above background. Note, we were unable to detect tethering at a 5 BoxB tethering cassette (data not shown). Thus, there is some unknown threshold of tethered Ago2 protein that is required for us to detect the signal. While the threshold is probably below 15 Ago2 or β-gal proteins (given the wide range in single mRNA tethering intensities we saw), a bright TnT biosensor is nonetheless a very large multi-protein/RNA complex whose size could interfere with or alter underlying biological processes. For example, the prevalent clustering we observed may not occur with such high probability when just one or two Ago2 proteins are recruited to an endogenous mRNA target. In the future it will therefore be important to reduce the number of stem loops required for detection, or perhaps develop a different tethering strategy with a smaller footprint. One promising strategy, for example, would be the use of fluorescently conjugated miRNA rather than EGFP-Ago2. The Walters lab demonstrated individual miRNA can be tracked in living cells (Pitchiaya et al., 2012, 2019, 2019), so in principle their method could be combined with ours to visualize both miRISC and target mRNA dynamics in a more natural setting. Finally, although our TnT biosensor associates with Ago2 independent of its interaction with a miRNA, it is likely that the complex does indeed contain a miRNA and may engage endogenous mRNA targets while tethered to the reporter. It is not clear what impact this might have on its functionality, although given the multivalent nature of Ago2 interactions with the miRISC machinery, it may be that even under natural conditions Ago2 interacts, albeit indirectly, with multiple mRNAs.

Moving beyond Ago2, the TnT biosensor can now easily be adapted to tether other proteins of interest. Indeed, tethering has frequently been used in the literature to study a wide variety of RNA binding proteins in diverse biological settings. For example, tethering has been used to investigate nonsense-mediated decay (Hickey et al., 2020), to screen for effects caused by RNA binding proteins (Luo et al., 2020), and to study specific protein domains (Fabian et al., 2011; Liu, Valencia-Sanchez, et al., 2005; Piao et al., 2010) or specific amino acid modifications (Golden et al., 2017). With the TnT biosensor, these studies can be expanded to the single-molecule level, where their impact can be more thoroughly investigated with higher spatiotemporal resolution. At the same time, in light of the presumed relocalization of the TnT biosensor to P-bodies upon Ago2 tethering, we believe our approach will be generally useful in targeting a biosensor to a specific subcellular environment, such as the nucleus, the ER, or mitochondria, where local translation can be studied (Buxbaum et al., 2015; Voigt et al., 2017). Finally, the TnT biosensor could be coupled with optogenetic dimerization domains to enable precision tethering with high spatiotemporal control (Benedetti et al., 2020; Kawano et al., 2015; Spiltoir & Tucker, 2019; Taslimi et al., 2014). In short, we anticipate the TnT biosensor will be a valuable new tool in the microscopy toolbox to investigate broad protein-mRNA interactions with unprecedented spatiotemporal resolution.

## Acknowledgements

We would like to thank all members of the Stasevich and Montgomery labs for innumerable critical conversations that helped develop and progress this project. We thank Dr. Hotaka Kobayashi and Dr. Rob Singer for sharing their preliminary data. We thank Dr. Ramesh Pillai for generously sharing with us the BoxB and λN plasmids. We thank Dr. Luis Aguilera for graciously supplying his code for bead alignment correction in Python. We thank Dr. O’Neil Wiggan and Nick Flint for helping with assay development for the fixed cell experiments. The JF646-HaloTag ligand was a generous gift from Dr. Luke Lavis at HHMI Janelia. This work was supported by grants from the National Institutes of Health (R35GM119728 to T.J.S and R35GM119775 to T.A.M). C.A.C. was also supported by the National Science Foundation NRT award DGE-1450032.

## Contributions

C.A.C. performed all experiments. C.A.C and T.J.S analyzed all data. C.A.C., T.J.S. and T.A.M. designed and planned all experiments. T.J.S. and T.M. assisted C.A.C. with microscopy and analysis and wrote the relevant methods sections. T.J.S., T.A.M. and C.A.C. wrote the main manuscript. C.A.C., T.M., T.A.M. and T.J.S. edited the manuscript. T.A.M. and T.J.S. acquired funding.

## Conflicts of Interest Statement

Nothing declared.

## Materials and Methods

### Plasmid construction

All expression vectors were designed using SnapGene software. The TnT biosensor construct (smFLAG-KDM5B-15xBoxB-24xMS2) was cloned using the smFLAG-KDM5B-24xMS2 plasmid from (Morisaki et al., 2016) and the pRL-5xBoxB construct from (Pillai et al., 2004). First, using restriction cloning, 5x BoxB repeats were copied from the pRL plasmid with AgeI sites on each end using PCR and inserted into the KDM5B reporter plasmid’s 3’ UTR at AgeI. Next, two 5x BoxB repeats were isolated from the pRL plasmid and added at the XbaI site, creating the final construct smFLAG-KDM5B-15xBoxB-24xMS2. The orientation of inserted BoxB stem loops was confirmed by Sanger sequencing (Quintara Biosciences). The plasmids were purified via midi-prep (Machery-Nagel) before loading.

The tetherable Ago2 (λN-EGFP-Ago2), non-tetherable Ago2 (EGFP-Ago2), and tetherable β-gal (λN-EGFP-β-gal) constructs were made by first swapping HA for EGFP in the plasmids λN-HA-hAgo2 or λN-HA-β-gal from (Pillai et al., 2004) using isothermal assembly. The EGFP sequence was obtained from the EGFP-hAgo2 plasmid (Addgene plasmid # 21981), which was a gift from Dr. Phil Sharp (Leung et al., 2006). Next, the entire open reading frame was inserted into the plasmid backbone from pFN24A HaloTag CMV*d3* Flexi Vector (Promega) so that the ORF was under a CMV-d3 minimal expression promoter. Lastly, we verified the plasmid sequence using whole-plasmid sequencing via de novo assembly (Gallegos et al., 2020) (Massachusetts General Hospital DNA sequencing core).

The sfGFP-H2B reporters were constructed from the plasmid sfGFP-H2B-C-10 (Addgene 56367, which was a gift from Dr. Michael Davidson) by inserting a UTR sequence at MfeI and HpaI in its 3’UTR. The “MRE” insert was created from a gene block containing the nucleotides 256-709 of the POLR3G 3’UTR, which was predicted to contain three miR-26-5p MREs with 50 nucleotides of endogenous UTR context flanking the first and last MRE. The mutant “mut MREs” insert additionally had two nucleotide mutations for each MRE (sequence and MRE locations obtained from TargetScan (*TargetScanHuman 7.2*, n.d.)). The insert’s orientation was verified by Sanger sequencing (Quintara Biosciences). The MRE-containing translation reporters used SpaghettiMonsterHA or SpaghettiMonsterFLAG constructs (sm**FLAG**-KDM5B-24xMS2 or sm**HA**-KDM5B-24xMS2 from (Morisaki et al., 2016)). The MRE or mutant POLR3G 3’UTR gene blocks were inserted at AgeI and XmaI sites in the 3’UTR of each FLAG and HA reporter. The insert orientation was verified by Sanger sequencing (Quintara Biosciences).

### Fab/Frankenbody Generation and MCP Purification

Cy3 α-FLAG-Fab were generated and affinity purified as previously described (Morisaki et al., 2016). Briefly, Fab were generated from monoclonal α-FLAG antibodies (Wako, 012-22384 Anti DYKDDDDK mouse IgG2b) using the Pierce Mouse IgG1 Fab and F(ab’)2 Preparation Kit (ThermoFisher). First, α-FLAG antibody were digested into Fab in a Zeba Desalt Spin Column (ThermoFisher) containing immobilized Ficin undergoing gentle rotation for 3-5 hours at 37°C. Fab were purified from the digest by centrifugation in a NAb Protein A column. Eluted Fab were concentrated to about 1 mg/mL and either conjugated with Cy3 or stored at 4°C. Cy3 labeling was performed in small batches of 100 μg Fab at a time. The dye was Cy3 *N*-hydroxysuccinimide ester (Invitrogen) dissolved in DMSO and either used immediately or stored at −20°C. For labeling, 100 μg of Fab was dissolved in a final volume of 100 μL of 100 mM NaHCO3 (pH 8.5) plus 1.33 μl of Cy3 dye. Fabs were labeled for about 2 hours at room temperature during gentle rotation and agitation. The Fab were separated from unconjugated dye in an equilibrated PD MiniTrap G-25 desalting column (GE Healthcare). Fab were concentrated in an Amicon Ultra-0.5 Centrifugal Filter Unit (NMWL 10 kDa; Millipore) to about 1 mg ml^−1^. The degree of labeling (*DOL*) was calculated using the following equation:

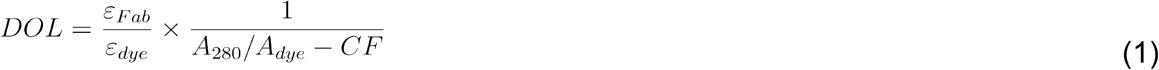

where ε_Fab_ is the extinction coefficient of Fab (70,000 M^−1^ cm^−1^), ε_dye_ is the extinction coefficient of the dye used for conjugation (150,000 M^−1^ cm^−1^ for Cy3), *A*_280_ and *A*_dye_ are the measured absorbances of dye-conjugated Fab fragments at 280 nm and at the peak of the emission spectrum of the dye (570 nm for Cy3), respectively, and *CF* is the correction factor of the dye (the ratio of the absorbances of the dye alone at 280 nm to at the peak; 0.08 for Cy3). If the DOL was less than 0.8, this protocol was repeated on the same Fab to increase their DOL to approximately 1. Fab were stored at 4°C.

MCP was generated as previously described (Morisaki et al., 2016). Briefly, MCP (His-HaloTag-2xMCP) was purified using its histidine tag with Ni-NTA Agarose (Qiagen) following the manufacturer’s instructions with minor modifications. The bacteria were lysed in a PBS-based buffer containing a complete set of protease inhibitors (Roche). In a gravity-flow column, binding to the Ni-NTA resin was carried out in the presence of 10 mM imidazole. After washing with 20 and 50 mM imidazole in PBS, the protein was eluted with 300 mM imidazole in PBS. The eluted protein was dialyzed against a HEPES-based buffer (10% glycerol, 25 mM HEPES pH 7.9, 12.5 mM MgCl_2_, 100 mM KCl, 0.1 mM EDTA, 0.01% NP-40 detergent, and 1 mM DTT) and stored at −80°C after snap-freezing by liquid nitrogen.

The GFP-tagged α-HA Frankenbody was generated and purified as previously described (Zhao et al., 2019). Briefly, α-HA Frankenbody was expressed in *E. coli* BL21 (DE3) pLysS cells transformed with pET23b-FB-GFP (Addgene plasmid 129593), induced with IPTG at O.D. ~0.6, and expressed at 18°C overnight. The protein was purified in two steps, first with HisTrap HP 5 mL columns (GE Healthcare) and next with a size-exclusion HiLoad Superdex 200 PG column (GE healthcare) in a HEPES-based buffer (25 mM HEPES pH 7.9, 12.5 mM MgCl_2_, 100 mM KCl, 0.1 mM EDTA, 0.01% NP40, 10% glycerol, and 1 mM DTT). Eluted α-HA Frankenbody was flash-frozen in liquid nitrogen and stored at −80°C. Only α-HA Frankenbody and MCP with 1-2 freeze-thaws were used in experiments.

### Cell culture

Human U2OS osteosarcoma cells were grown in 10% (v/v) FBS (Atlas) media DMEM (Thermo Scientific) supplemented with 1 mM L-glutamine (Gibco) and 1% (v/v) Pen/Strep (Invitrogen/Gibco) and grown at 37°C in 5% CO_2_. Cell density was maintained between 20-80%. U2OS cells were purchased from ATCC and were authenticated by STR profiling by ATCC and morphological assessments. We also confirmed that the cell line tested negative for mycoplasma contamination.

### Bead Loading

To bead load TnT plasmid and protein components, cells were plated at 70% confluency on 35 mm glass bottom chambers (MatTek) 1-2 days before imaging. Cells were bead loaded as described previously (Hayashi-Takanaka et al., 2011; McNeil & Warder, 1987). Briefly, the media was changed to 10% FBS Opti-MEM (Thermo Scientific) before bead loading. The components were mixed together in a total volume of 4-8 μL: ~100 μg/mL of Cy3 α-FLAG Fab, ~33 μg/ml of purified HaloTag-2xMCP protein, plasmids, and 1x PBS if needed to fill to volume. Concentrations of 1.5 or 1.75 μg of DNA were used for the GFP-containing plasmids (either λN-EGFP-Ago2, λN-EGFP-β-gal, or EGFP-Ago2) or the reporter plasmid (pUb-smFLAG-KDM5B −15xBoxB-24xMS2), respectively. After removing the media, the 4 μL mixture of protein and DNA were pipetted on top of the cells, followed by sprinkling a monolayer of ~106 μm glass beads on top of the cells (Sigma Aldrich). The chamber was tapped ~20 times gently, then the media were replaced. After ~2.5 hours, cells were washed three times in phenol-free DMEM with 10% FBS and 1 mM L-glutamine. Before imaging, cells were stained with 200 nM JF646 HaloTag ligand for 15 minutes, followed by three washes in DMEM with 10% FBS and 1 mM L-glutamine. Cells were moved to the microscope stage-top incubator for imaging ~3 h post bead loading.

### Transfection

Transfection was performed in the fixed cell experiments where no protein loading was needed. All transfections were performed with the LTX Lipofectamine with Plus Reagent kit (Thermo Fisher), per the manufacturer’s instructions. Briefly, cells were washed and the media was replaced with 1.75 mL Opti-MEM (Thermo Scientific) directly before transfection. The transfection solution included 2.5 μg DNA plasmid, 7.5 μL Plus reagent, and 7.5 μL Lipofectamine and Opti-MEM brought the solution to 250 μL. This solution was incubated for 5-15 minutes at room temperature before being added to the cell chamber. Cells were incubated in this transfection solution for 2-4 hours before the media was changed back to 10% FBS-DMEM. For the TnT biosensor, 1.5 μg smFLAG-KDM5B-15xBoxB-24xMS2 or smFLAG-KDM5B-24xMS2 plasmid plus μg of either λN-EGFP-Ago2, λN-EGFP-β-gal, or EGFP-Ago2 plasmid were used. For sfGFP-H2B reporter assays, 2.5 μg plasmid was used.

### Cell Imaging with the confocal microscope

Fixed-cell images were acquired on an Olympus (IX83) Inverted Spinning Disk Confocal Microscope with a cascade II EMCCD camera. The objectives used were the 40x oil-immersion (0.24 μm per pixel) and the 100x oil-immersion (0.096 μm per pixel). For smiFISH experiments, the 405 nm, 488 nm, 561 nm, and 647 nm lasers were used. Each location captured a 15 image Z-stack with a step size of 0.35 μm which was max-projected. For the sfGFP-H2B reporter assay, the 405 nm, 488 nm lasers were used. Each location captured a 15 image Z-stack with a step size of 0.35 μm which was max-projected.

### Cell Imaging with the HILO microscope

All live-cell imaging was performed on a custom built widefield fluorescence microscope with a highly inclined thin illumination scheme (Tokunaga et al., 2008)) described previously (Morisaki et al., 2016). Briefly, the microscope equips three solid-state laser lines (488, 561, and 637 nm from Vortran) for excitation, an objective lens (60X, NA 1.49 oil immersion, Olympus), an emission image splitter (T660lpxr, ultra-flat imaging grade, Chroma), and two EMCCD cameras (iXon Ultra 888, Andor). Achromatic doublet lenses with 300 mm focal length (AC254-300-A-ML, Thorlabs) were used to focus images onto the camera chips instead of the regular 180 mm Olympus tube lens to satisfy Nyquist sampling (this lens combination produces 100X images with 130 nm/pixel). The far-red signal of mRNA visualized by JF646 HaloTag MCP was imaged on one camera, and the red signal of translation visualized by Cy3 α-FLAG-Fab and the green signal of GFP-tagged Ago2, β-gal, or α-HA Frankenbody were imaged on the other camera. A high-speed filter wheel (HS-625 HSFW TTL, Finger Lakes Instrumentation) was placed in front of the second camera to minimize the bleed-through between the red and the green signals (593/46 nm BrightLine for the red and 510/42 nm BrightLine for the green, Semrock). The focus was maintained throughout the experiments with the CRISP Autofocus System (CRISP-890, Applied Scientific Instrumentation). The Z-stack images were taken with a piezoelectric stage (PZU-2150, Applied Scientific Instrumentation). The laser emission, the camera integration, the piezoelectric stage movement, and the emission filter wheel position change were synchronized by an Arduino Mega board (Arduino). Image acquisition was performed using opensource Micro-Manager software (1.4.22) (Edelstein et al., 2010).

Live cells were placed into a stage-top environmental chamber at 37°C and 5% CO_2_ (Okolab) to equilibrate for at least 30 minutes before image acquisition. Imaging size, exposure time, and the vertical shift speed was set to 512 x 512 pixels^2^, 53.64 msec, and 1.13 microsec, respectively. This resulted in the imaging rate of 13 Hz (70 msec per image). To capture the whole thickness of the cytoplasm of U2OS cells, 13 Z-stack of step size 500 nm were imaged such that the Z-position changed every 2 images (for the red and the green signals at each stack). This resulted in a maximal cellular imaging rate of 0.5 Hz (2 sec per volume). When needed, delays between volume captures were used to image at lower frame rates. For longterm single molecule tracking (**Fig. 4 and Sup. Fig. 4)**, photobleaching was minimized by imaging mRNA (JF646) every 10 seconds, while imaging tethering and translation (GFP and Cy3) every 100 seconds. Laser powers for all movies were: 15-20 mW for 637 nm, 9-20 mW for 488 nm and 5-15 mW for 561 nm with an ND10 neutral density filter at the beam expander.

TetraSpeck Fluorescent Microspheres (100 nm, ThermoFisher Scientific) mounted on a MatTek chamber were imaged at the end of each imaging session in order to correct for the slight shift in the alignment of the two cameras. These images of beads were used to generate a transformation matrix using either the GeometricTransformation function in Mathematica (Wolfram Research) or the ProjectiveTransform function in scikit-image in Python to correct for offsets in detected particle positions in each channel.

### Co-Immunofluorescence and RNA smiFISH

Single-Molecule Inexpensive Fluorescent In Situ Hybridization (smiFISH) plus Immunofluorescence (IF) experiments were performed as previously described (LGC Biosearch Technologies, n.d.; Tsanov et al., 2016). Briefly, cells were transfected (as described in the “**Transfection**” section). After 24 hours, cells were washed 3 times in PBS and fixed in 4% paraformaldehyde (Sigma Alrich) PBS for 20 minutes at 37°C. Cells were washed then permeabilized in 0.1 mM TritonX (Sigma Alrich) in PBS for 20 minutes.

Next, smiFISH was performed at room temperature unless otherwise specified. The smFISH hybridization and wash buffers were from Biosearch Technologies Stellaris buffers (SMF-HB1-10, SMF-WA1-60, and SMF-WB1-20) and used as stated in the Stellaris protocol for adherent cells. The probe hybridization was performed for ~12 hours at 37°C. The probe set for smFLAG (5’) was designed using the open-source R script Oligostan (Tsanov et al., 2016), and the probe set for MS2 (3’) was copied from (Tsanov et al., 2016).

Lastly, immunostaining was performed at room temperature unless otherwise specified. Cells were blocked with 1:4 diluted Blocking One-P (Nacalai Tesque) in PBS for 1 h, then stained with a 1:500 dilution of α-RCK antibody (MBL, PD009) in 1:4 diluted Blocking One-P for 1 h. Cells were washed for 30 minutes four times in 0.1% Tween-20 PBS, blocked as above for 1 h, then incubated in DyLight 405-AffiniPure F(ab’)2 α-Rabbit antibody (Jackson ImmunoResearch) diluted 1:2000 in 1:4 diluted blocking buffer. Cells were washed for 30 minutes four times in 0.1% Tween-20 PBS and mounted in ProLong Diamond AntiFade Mountant (Thermo Fisher). Following these procedures, fixed cells were imaged as described in “**Cell Imaging with the confocal microscope**”.

### Live-cell nuclear reporter accumulation assay

For experiments in **Fig. 2B,C** and **Sup. Fig. 2A**, cells were bead loaded with 0.5 μg of purified Cy3 α-FLAG Fab, 130 nm of purified HaloTag-2xMCP, 1.5 μg of the TnT biosensor plasmid (smFLAG-KDM5B-15xBoxB-24xMS2) and 1.0 μg of either tetherable Ago2 (λN-EGFP-Ago2), tetherable β-gal (λN-EGFP-β-gal), or non-tetherable Ago2 (EGFP-Ago2), as described in the “**Bead Loading**” section. The HaloTag was labeled for 15 minutes with JF646 HaloTag Ligand 1 hour after bead loading, after which cells were moved to the microscope stage-top incubator. At 4 hours post-bead loading, the microscope was programmed to visit multiple locations and take one full cell volume (13 Z-planes with a 0.5 μm step size) every 30 minutes. This continuous imaging was important to identify and exclude cells that died or divided, or moved away.

All videos were max-Z projected. Masks were hand-drawn on the 4 hour (early) and 16 hour (late) frames to isolate the nucleus and cytoplasm and used to measure the mean intensity of Cy3 α-FLAG Fab in these locations. Reporter accumulation in the nucleus was calculated:

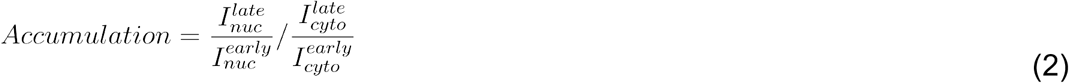

where 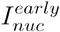 and 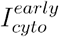 were the measured intensity of the nucleus or cytoplasm, respectively, at the 4 hour timepoint, while 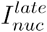 and 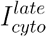 were the measured intensity of the nucleus or cytoplasm, respectively, at the 16 hour timepoint. Two of three experimental days were performed blinded.

### Fixed-cell TnT nuclear reporter accumulation assay

For experiments in (**Sup. Fig. 2B**), cells were transfected with the plasmid smFLAG-KDM5B-15xBoxB-24xMS2 and with either tetherable Ago2 (λN-EGFP-Ago2), tetherable β-gal (λN-EGFP-β-gal), or non-tetherable Ago2 (EGFP-Ago2), as described in the “**Transfection**” section above. After 24 hours, cells were fixed in 4% paraformaldehyde-PBS for 20 minutes at 37°C then permeabilized in 0.1% TritonX100 for 20 minutes at 37°C.

Cells were blocked with 1:4 diluted Blocking One-P (Nacalai Tesque) in PBS for 1 h, then stained with 0.5 μg Cy3 α-FLAG Fab diluted to 100 μL in 1:4 diluted Blocking One-P. Cells were washed for 30 minutes four times in PBS and imaged directly, as described in the “**Cell Imaging with the HILO microscope**” section. Any images with multiple cells were cropped such that in the end each image showed a single cell, and a mask was hand-drawn around each cell. The mean Fab signal in the masked cell was measured per image. The imaging and analysis of all three experiments were performed blinded.

### Fixed-cell miRNA target site reporter nuclear accumulation assay

For experiments in (**Sup. Fig. 5B**), cells were transfected with 2.5 ug of the plasmid sfGFP-H2B-MRE-POLR3G-3’UTR or sfGFP-H2B-mut-POLR3G-3’UTR, as described in the “**Transfection**” section above. After 24 hours, cells were fixed in 4% paraformaldehyde-PBS for 20 minutes at 37°C then stained in 1:2000 diluted Hoechst 33232 (ThermoFisher) for 10 mins.

Image analysis was performed using custom code in Mathematica (Wolfram Research) to detect and measure the fluorescence intensities of all nuclei. Each background subtracted image in the Hoechst channel was binarized (using a sensible threshold) to create a nuclear mask. Using Mathematica’s built-in function “ComponentMeasurements,” nuclei that crossed the image edge were discarded, and masks were shape- and size-selected so that only whole, single nuclei were detected. The mask was applied to the 488 nm channel of the image and the total sfGFP fluorescence signal of each nucleus was measured. The imaging and analysis of all three experiments were performed blinded. All code is available on Github.

### MRE-reporter assay

This experiment (**Fig. 5, Sup. Fig. 5C**) used a modified TnT reporter using endogenous MREs instead of BoxB tethering cassette. Using TargetScan (*TargetScanHuman 7.2*, n.d.), we chose a stretch of the POLR3G 3’UTR that had three miR-26-5p sites targeted by miRNA that are abundant in U2OS cells (Jerez et al., 2019). As a control (called the MUT reporter), we mutated two nucleotides in each miR-26-5p site, rendering them untargetable by Ago2 (Mayr et al., 2007) (plasmids are further described above in “**Plasmid construction**”). Each experiment used two cell chambers. Cells were loaded with the plasmids described below, along with 0.5 μg of Cy3 α-FLAG Fab, 0.5 μg of GFP-tagged α-HA Frankenbody, and 130 nm of HaloTag-2xMCP (further description of loading in the section “**Bead Loading**”). Each replicate experiment used one chamber loaded with 1 μg of smFLAG-KDM5B-MRE-POLR3G-3’UTR-24xMS2 and 1 μg of smHA-KDM5B-mut-POLR3G-3’UTR-24xMS2 and a second chamber loaded with the reversed combination of smHA-KDM5B-MRE-POLR3G-3’UTR-24xMS2 plus 1 μg of smFLAG-KDM5B-mutPOLR3G-3’UTR-24xMS2. Both chambers were then imaged on our custom HILO microscope. For each cell, 20 frames (11-13 Z-stacks with a 0.5 μm vertical step size) were imaged at a 2 frames*s^−1^ frame rate. Two of the three experiments were performed blinded.

### Puromycin translation inhibition assay

Puromycin assays (**Fig. 1D, Sup. Fig. 1B,C,E**) were performed as previously described (Morisaki et al., 2016). Briefly, cells were bead loaded with the TnT components along with either tetherable Ago2 (λN-EGFP-Ago2), tetherable β-gal (λN-EGFP-β-gal), or non-tetherable Ago2 (EGFP-Ago2). While on the microscope, cells were treated with 50 μg/mL of puromycin. Cells were imaged continuously for 5 frames before and up to 60 frames aftertreatment at a rate of one full cell volume (11-13 Z-planes with a 0.5 μm step size) per 10 or 30 seconds. All mRNA foci colocalized with a Cy3 α-FLAG Fab translation signal were detected by hand and further categorized by colocalized tethering signal. The count of tethered/untethered, translating mRNA per cell were quantified per frame.

### Single particle detection, tracking, and intensity measurement

Image processing was done in the software program Fiji (Schindelin et al., 2012) by Z-projecting each time frame of the 3-dimensional movie. These images were then analyzed using custom Mathematica code (Wolfram Research, available on GitHub) to detect single particles in a semi-automated fashion. Briefly, for each image the background was masked by a hand-drawn outline of the cell and each image was binarized using a bandpass filter to highlight particles between 1 and 7 pixels in size (96 nm/pixel for the confocal, 130 nm/pixel for our custom HILO microscope). The mRNA channel was used to detect spots agnostic to tethering or translation status. Mathematica’s built-in “ComponentsMeasurements” function was used to select and filter out larger aggregates from single particles and to categorize mRNA into groups based on the presence of detectable particles with signals above background in each channel. When needed, particles were tracked. Briefly, detected mRNA were linked to the closest mRNA in the next time frame, with a maximum step size of 10 pixels and a shortest track length of 5 frames. All detected particles and tracks were confirmed by hand when necessary.

To quantify particle intensities in each channel, intensities were local-background subtracted. This involved three steps using a 15 x 15 pixel crop (130 nm/pixel) of each detected particle: (1) the central signal was calculated as the mean intensity in a centered disc of diameter 3 pixels; (2) the background signal was quantified by measuring the median intensity in four 9-pixel quadrants in the corners of the 15 x 15 pixel crop; (3) The particle’s signal intensity in each channel was computed by subtracting the background signal from the central signal. For fixed cell images using the confocal microscope (**Fig. 3C,D and Sup. Fig. 3**), steps 1 and 2 were modified such that the central disc was 5 pixels in diameter to accommodate larger P-bodies (96 nm/pixel), and the background was the mean intensity in an outer ring of diameter 15 pixels and width 2 pixels (96 nm/pixel).

For the long-term, single-molecule tracking experiments in **Fig. 4 and Sup. Fig 4**, intensity measurements were further refined to get the highest possible signal to noise. For this, we used custom Python code to do “best-Z” projections rather than max-Z projections. This was achieved using the positions of particles tracked in 2D to make a set of 3D crops around each detected particle at each timepoint. Each 3D crop had an XYZ dimension of 11 x 11 x 3 voxels, respectively, where each voxel had an XYZ dimension of 130 nm x 130 nm x 500 nm, respectively. In these 3D crops, the 3 chosen Z-slices were Z_best_ ± 1 such that Z_best_ was the Z-slice having a maximal mean mRNA intensity inside a central disc of diameter 6 pixels (130 nm/pixel). Four steps were then used to quantify the signal intensities in each channel from individual 3D crops: (1) the 3 best Z slices were max-Z projected (3-frame moving averages of these best-Z projected crops are displayed in **Fig. 4 and Sup. Fig. 4**; the only exception is **Fig. 4A**, where every frame is shown); (2) the central signal was calculated as the mean intensity in a centered disc of diameter 6 pixels (130 nm/pixel); (3) The background signal was calculated as the mean intensity in a surrounding ring of diameter 10 pixels and width 1 pixel (130 nm/pixel); and (4) The signal intensity was computed by subtracting the background signal from the central signal. To aid in this analysis, each step was viewed using Napari (napari contributors, 2019). From the measured signal intensities, CSV files were generated and Seaborn (Waskom, 2020) was used to create scatter plots and intensity vs. time plots. All Python code for this analysis is available on Github.

### Harringtonine ribosome runoff assay

To generate Harringtonine (HT) runoff curves, cells were bead loaded with the TnT components along with either tetherable Ago2 (λN-EGFP-Ago2), tetherable β-gal (λN-EGFP-β-gal), or non-tetherable Ago2 (EGFP-Ago2). While on the microscope, cells were treated with 3 μg/mL of Harringtonine (Cayman Chemical) to inhibit translation initiation (Huang, 1975). Cells were continuously imaged for 5 minutes before and 60 minutes after addition of HT, at a rate of one full cell volume (11-13 Z-planes with a 0.5 μm step size) per minute. For photobleach correction, a hand-drawn mask was used to find the mean, background subtracted whole-cell intensity through time (**Sup. Fig. 4C**). The single exponential fit of the signal decay was calculated per channel and divided from single mRNA translation intensities through time. Images were preprocessed and all mRNA in cells were detected as described in the “**Single particle detection, tracking, and intensity measurement**” section. Intensities of particles in 3D were quantified using best-Z projections rather than max-Z projections. From these, we plotted the average single-molecule signals from single cells treated with HT through time (with the corresponding average best-Z projected crops shown above in **Fig. 4G and Sup. Fig. 4B**). The average of the first four timepoints was used to normalize single-cell runoff curves so they began at a value of one.

Each runoff curve was fit to a biphasic model. In the model, a fraction *f* of ribosomes run off transcripts with an elongation rate *v*1 and the remaining fraction (1 − *f*)run off transcripts with an elongation rate *v*2. The full model as a function of time t can be written as:

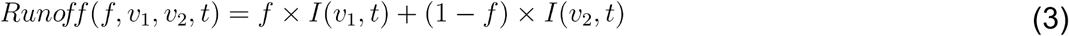

Here *I*(*v, t*)describes a monophasic ribosome runoff curve in which all ribosomes are assumed to run off the transcript with an average elongation rate of *v* (Pichon et al., 2016):

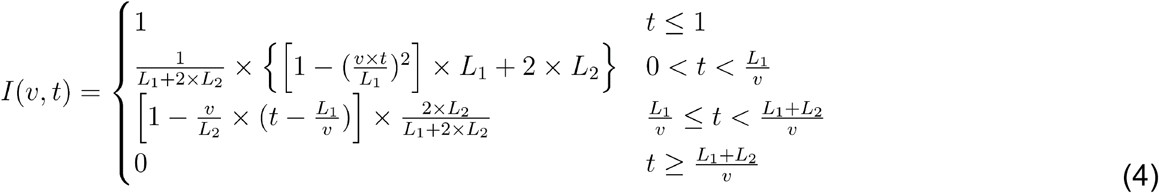

where *L*1 is the length of the tagged portion of the ORF in our TnT reporter (336 aa) and *L*2 is the length of the untagged portion (1549 aa). After fitting, the runoff halftime *t*_1/2_ was determined by setting the fitted runoff curve to 0.5. Fitting was performed with Python using the command scipy.optimize.curve_fit from SciPy. All code is available on Github.

## Data availability

The raw data that was used to generate Fig. 1C, Sup. Fig. 1B,C,E, Fig. 2D,E, Sup. Fig. 2B-D, Fig. 3B,D, Sup. Fig. 3A,B, Fig. 4A,-H, Sup. Fig. 4A-G, Fig. 5C, and Sup. Fig. 5B,CA are stored in a publicly available repository: “Datasets associated with ‘Imaging translational control by Argonaute with single-molecule resolution in live cells’”. https://doi.org/10.6084/m9.figshare.c.5395800.v2 (Cialek, 2021). All other data supporting the findings of this study are available from the corresponding author on request.

## Source Data

Source Data for **Figure 1**

- Fraction of tethered mRNA per cell for **Fig. 1C**
- Count of translating mRNA per cell in puromycin translation inhibition experiments for **Sup. Fig. 1B,C,E**

Source Data for **Figure 2**

- Fraction of translating, tethered mRNA per cell for **Fig. 2D**
- Translation intensity of single mRNA for one experimental day for **Fig. 2E**
- Intensity of reporter in nucleus per cell (fixed) **Sup. Fig. 2B**
- Count of mRNA per cell classified on tethering and translation status for **Sup. Fig. 2C**
- Translation intensity of single mRNA for three experimental days for **Sup. Fig. 2D**

Source Data for **Figure 3**

- Intensity of mRNA puncta over 12 hours for **Fig. 3B**
- Fraction of mRNA in P bodies per cell for **Fig. 3D**
- Fraction of intact mRNA per cell and in/out of P-bodies for **Sup. Fig. 3A**
- Intensity ratio of 3’ / 5’ per mRNA in vs out of P-bodies in cells expressing tetherable Ago2 for **Sup. Fig. 3B**

Source Data for **Figure 4**

- Raw intensity of translation, Ago2 or β-gal, and mRNA for single mRNA tracks for **Fig. 4A,B**
- Double-renormalized intensity of translation, Ago2 or β-gal, and mRNA for single mRNA tracks for **Fig. 4C,D,E,F** and **Sup. Fig. 4A**
- Translation intensity per mRNA for Harringtonine runoff assay for **Fig. 4G** and **Sup. Fig. 4B,D**
- Whole-cell intensity during Harringtonine runoff assay for photobleach correction measurement for **Sup. Fig. 4C**
- Slow rate, fast rate, and fast fraction data for each cell for Harringtonine runoff assay for **Sup. Fig. 4E,F,G**
- Translation half-times for each cell for Harringtonine runoff assay for **Fig. 4H**

Source Data for **Figure 5**

- Intensity of FLAG and HA per mRNA for miRNA site targeting translation assay for **Fig. 5C** and **Sup. Fig. 5C**
- sfGFP intensity per nucleus for miRNA site targeting translation assay for **Sup. Fig. 5B**

## Supplemental Video Legends

All supplemental videos are stored in a publicly available repository: “Datasets associated with ‘Imaging translational control by Argonaute with single-molecule resolution in live cells’”. https://doi.org/10.6084/m9.figshare.c.5395800.v2 (Cialek, 2021).

**Supplemental Video 1. Simultaneous imaging of single-molecule translation and tethering with the TnT biosensor.** A video showing a representative cell loaded with the TnT components (plasmids encoding λN-EGFP-Ago2, smFLAG-KDM5B-15xBoxB-24xMS2 mRNA reporter, Cy3-FLAG-Fab, and JF646 HaloTag MCP). The video was acquired at 0.5 frames/second for 40 seconds total (blue, tetherable Ago2; green, translation; red, mRNA). The circles mark single mRNA being (i) translated and Ago2-tethered, (ii) translated and Ago2-untethered, or (iii) non-translated and Ago2-tethered. Scale bar, 10 μm.

**Supplemental Video 2. TnT biosensor translation signals disappear upon puromycin treatment.** A video showing a representative cell loaded with the TnT components (plasmids encoding λN-EGFP-Ago2, smFLAG-KDM5B-15xBoxB-24xMS2 mRNA reporter, Cy3-FLAG-Fab, and JF646-stained MCP) and treated with puromycin. The video was acquired at 0.1 frames/second for 30.67 minutes total (blue, tetherable Ago2; green, translation; red, mRNA). 50 mg/ml puromycin was added at Time = 0 (frame 5). Scale bar, 10 μm.

**Supplemental Video 3. Translationally-silenced TnT biosensors cluster and coalesce with other mRNA foci over hours.** A video showing a translationally-silenced, Ago2-tethered TnT biosensor clustering and coalescing with other mRNA foci over hours. The video comes from a representative cell 4 - 16.5 hours after loading the TnT components (plasmids encoding λN-EGFP-Ago2 and smFLAG-KDM5B-15xBoxB-24xMS2 mRNA reporter, Cy3-FLAG-Fab, and JF646 HaloTag MCP). The cell was imaged every 30 minutes for 12 hours (blue, tetherable Ago2; green, translation; red, mRNA). Scale bar, 10 μm.

**Supplemental Video 4. Ago2 tethering silences translation gradually at a single mRNA.** A video showing a sample track of a single TnT biosensor in cells after loading TnT components (plasmids encoding λN-EGFP-Ago2, smFLAG-KDM5B-15xBoxB-24xMS2 mRNA reporter, Cy3-FLAG-Fab, and JF646 HaloTag MCP). The video was acquired at 0.1 frames/second in the mRNA channel and 0.01 frames/second in the translation (green) and Ago2 tethering (blue) channels for a total of 75 minutes. The frame size is 0.6 μm x 0.6 μm.

**Supplemental Video 5. Long-term silencing of Ago2-tethered mRNA.** A video showing a sample track of a single TnT biosensor in cells after loading TnT components (plasmids encoding λN-EGFP-Ago2, smFLAG-KDM5B-15xBoxB-24xMS2 mRNA reporter, Cy3-FLAG-Fab, and JF646 HaloTag MCP). The video was acquired at 0.88 frames/second in the mRNA channel and 0.088 frames/second in the translation (green) and Ago2 tethering (blue) channels for a total of 79.2 minutes. The frame size is 0.46 μm x 0.46 μm.

**Supplemental Video 6. Ago2-tethered mRNA loses translation signal in a rare splitting event at a single mRNA.** A video showing a sample track of a single TnT biosensor in cells after loading TnT components (plasmids encoding λN-EGFP-Ago2, smFLAG-KDM5B-15xBoxB-24xMS2 mRNA reporter, Cy3-FLAG-Fab, and JF646 HaloTag MCP). The video was acquired at 0.1 frames/second in the mRNA channel and 0.01 frames/second in the translation (green) and Ago2 tethering (blue) channels for a total of 75 minutes. The frame size is 1.15 μm x 1.15 μm.

**Supplemental Video 7. Sample Harringtonine runoff experiment in cells expressing tetherable Ago2.** A video showing a sample cell after loading TnT components (plasmids encoding λN-EGFP-Ago2, smFLAG-KDM5B-15xBoxB-24xMS2 mRNA reporter, Cy3-FLAG-Fab, and JF646-stained MCP). Harringtonine was added at Time = 0 (frame 5). The video was acquired at 1 frames/minute for a total of 65 minutes (blue, tetherable Ago2; green/grayscale, translation; red, mRNA). Scale bar, 10 μm.

**Supplemental Video 8. Tracking the translation of single biosensors containing either endogenous and mutated MREs in the same cell.** A video of a sample cell expressing modified biosensors with either 10 FLAG epitopes and endogenous MREs or 10 HA epitopes and mutated MREs. The cells were loaded with these plasmids along with Cy3 α-FLAG Fab to label FLAG translation (green), GFP-tagged α-HA Frankenbody to label HA translation (blue), and JF646 HaloTag MCP to label mRNA (red). The video was acquired at 0.5 frames/second for a total of 40. Scale bar, 10 μm.

---

**Supplemental Figure 1.**
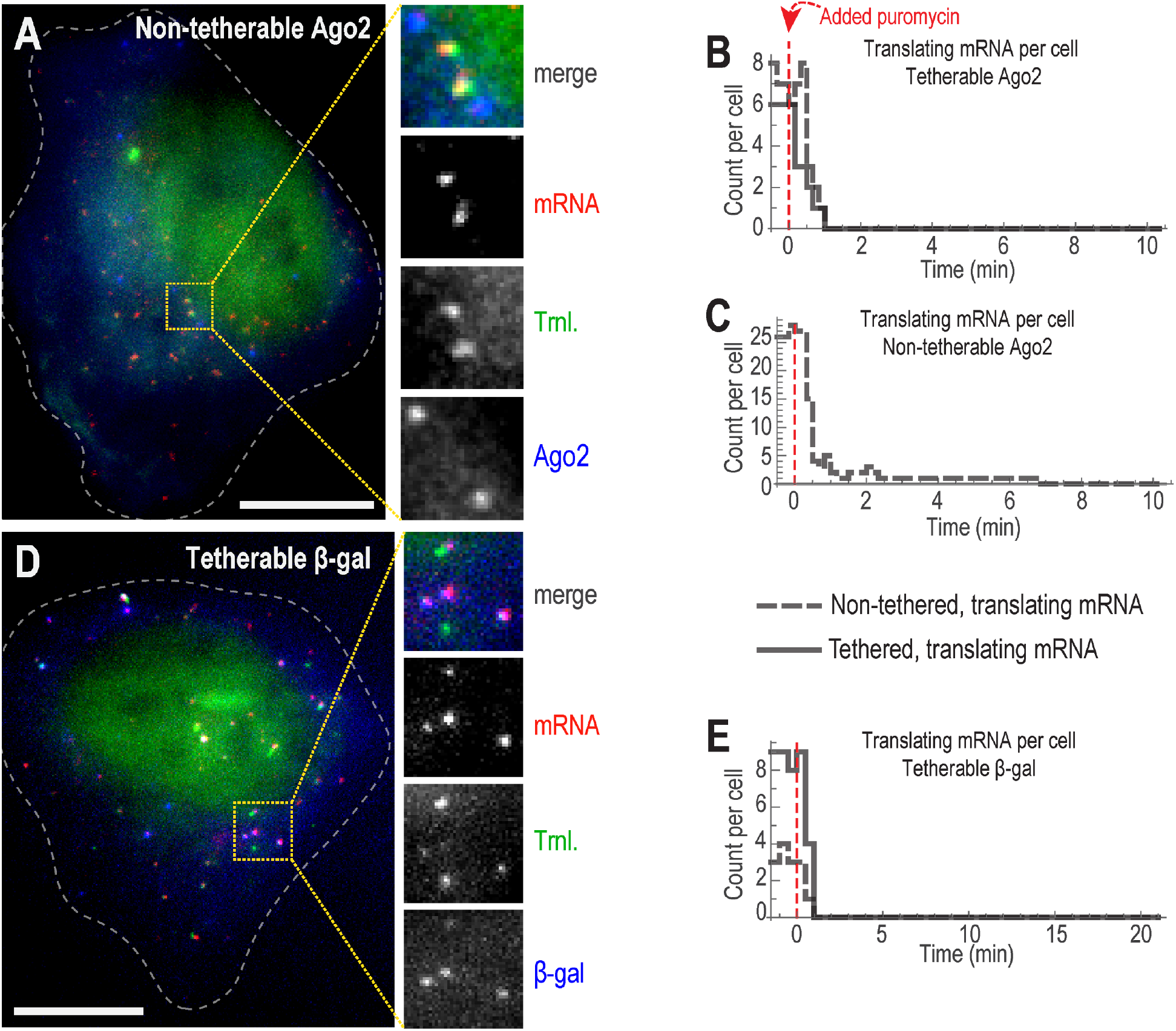
The TnT biosensor can be used to simultaneously track translation and tethering to reporter mRNA. **A** A representative cell expressing non-tetherable Ago2 (EGFP-Ago2). The image was acquired approximately 4 hours after loading the TnT components (smFLAG-KDM5B-15xBoxB-24xMS2 mRNA reporter, Cy3-FLAG-Fab, and JF646 HaloTag MCP). The dashed line marks the cell boundary. Scale bars, 10 μm. **B-C** Line plots showing the number of mRNA actively being translated in a single, TnT-loaded cell expressing tetherable Ago2 (B) or non-tetherable Ago2 (C), prepared as described above. Cells were treated with puromycin at Time = 0, as indicated by the dashed, vertical red line, and imaged for 10-20 minutes (min). Solid and dashed lines correspond to tethered and untethered mRNA, respectively. **D** A representative cell expressing tetherable β-gal (λN-EGFP-β-gal) and the TnT components. The image was acquired approximately 4 hours after loading the TnT components. The dashed line marks the cell boundary. Scale bar, 10 μm. **E** Same as B,C, but now for a single cell expressing tetherable β-gal.

**Supplemental Figure 2.**
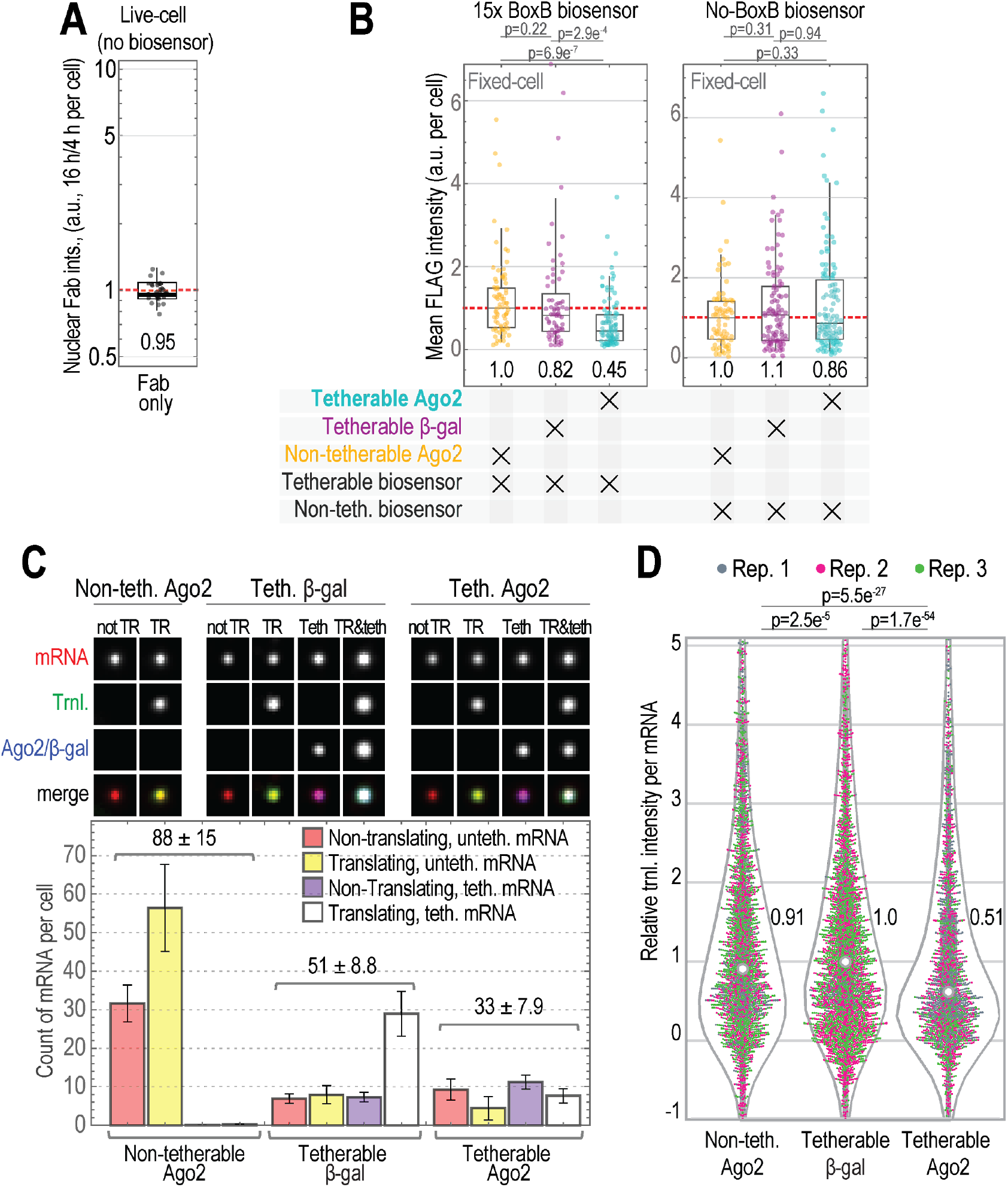
Impact of Ago2 tethering on translation. **A** Box plot displaying nuclear accumulation of Fab in the absence of the TnT biosensor. Each data point is the ratio of Fab nuclear intensity at 16 hours to 4 hours after Fab loading. **B** Box plot showing the mean intensity of the TnT reporter protein KDM5B (as marked by Fab) in fixed cells expressing the TnT biosensor (with or without the BoxB tethering cassette) as well as non-tetherable Ago2, tetherable β-gal, or tetherable Ago2. Cells were fixed 24 hours after TnT biosensor loading and stained with α-FLAG Fab. The mean cellular intensity was measured. N=77 non-tetherable, 67 B-gal-tetherable, and 97 Ago2-tetherable cells (left), and N=70 non-tetherable, 102 B-gal-tetherable, and 114 Ago2-tetherable cells (right). **C** Box plot of all mRNA per live, TnT-loaded cells from **Fig. 2E** (1 of 3 replicates). The mRNA were categorized based on the presence or absence of detectable translation (TR) and/or Ago2/β-gal tethering (Teth) signals. Above, representative background-subtracted and auto-adjusted mean crops from each category (18 x 18 pixels^2^; 130 nm/pixel). Note, because very few non-tetherable Ago2 cells had tethering signals, crops showing tethering in this case are not shown. Below, mean counts per cell for each category (color-coded bars). The total number of mRNA particles per cell were N_total_= 88 ± 15 (mean±SEM), 51 ± 8.8, and 33 ± 7.9 for non-tetherable Ago2, tetherable β-gal and tetherable Ago2, respectively. Error bars show SEM. N= 18 (1513), 28 (1158), 18 (521) cells (mRNA) for non-tetherable Ago2, tetherable β-gal, and tetherable Ago2, respectively. **D** Violin plot showing all experimental replicates (discernable by marker color) from the experiment in **Fig. 2E**. P values were calculated using the Bonferroni-corrected Mann-Whitney test. N = 55 (3,153), 39 (4,947) and 52 (3,959) cells (mRNA) for non-tetherable Ago2, tetherable β-gal, and tetherable Ago2, respectively.

**Supplemental Figure 3.**
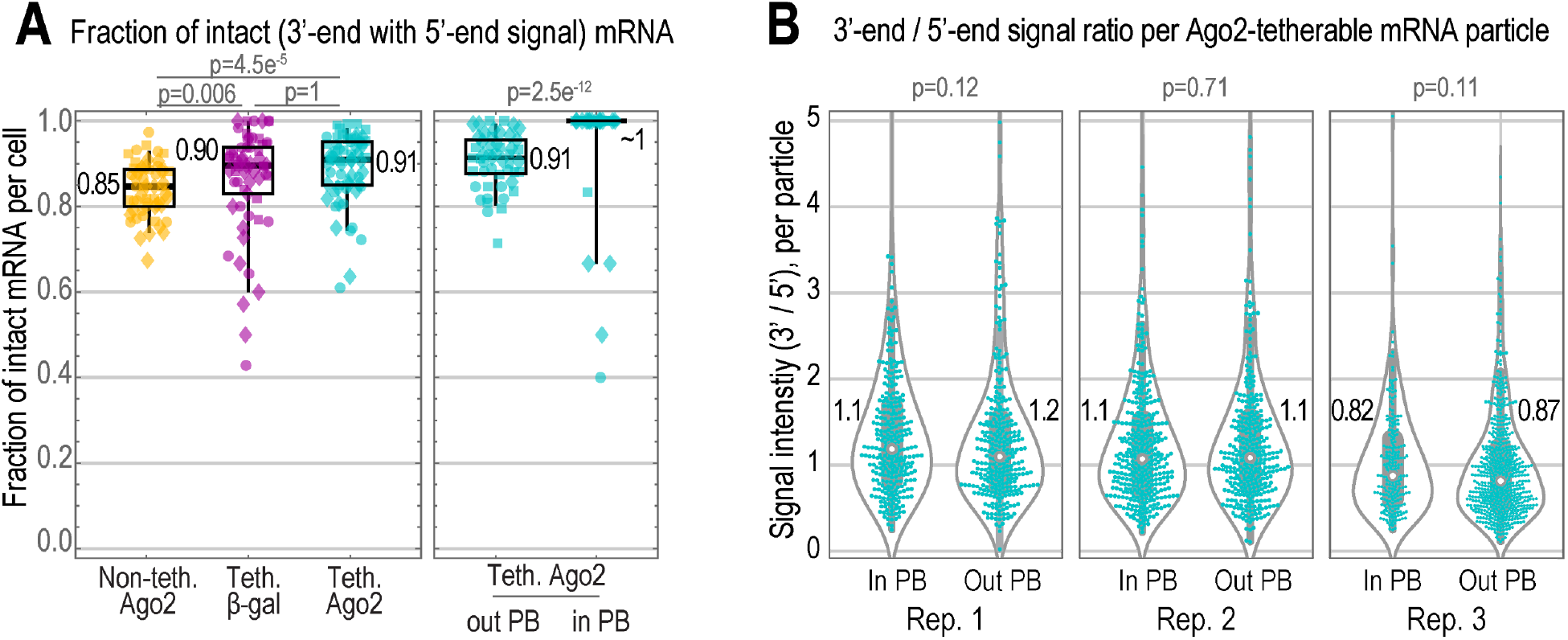
Most TnT biosensors in cells have 5’ and 3’ smiFISH signals. These data come from the experiment shown in **Fig. 3C-D**. **A** Box plots quantifying the fraction of mRNA per cell that were detected by their 3’ smiFISH signal and also had a 5’ smiFISH signal (yellow, non-tetherable Ago2; purple, tetherable β-gal; cyan, tetherable Ago2; different shapes mark 1 of 3 replicate experiments). Left, tetherable Ago2 versus controls. Right, tetherable Ago2 signals in P-bodies (PBs) versus out. P values were calculated from Mann-Whitney tests (A, left was Bonferroni-corrected). **B** Box plots showing the ratio of 3’ to 5’ smiFISH signals per each mRNA focus (including multi-mRNA clusters) in or out of PBs in all cells expressing tetherable Ago2 over three experimental replicates. The median ratio value per replicate experiment was calculated. P values were calculated from Mann-Whitney tests.

**Supplemental Figure 4.**
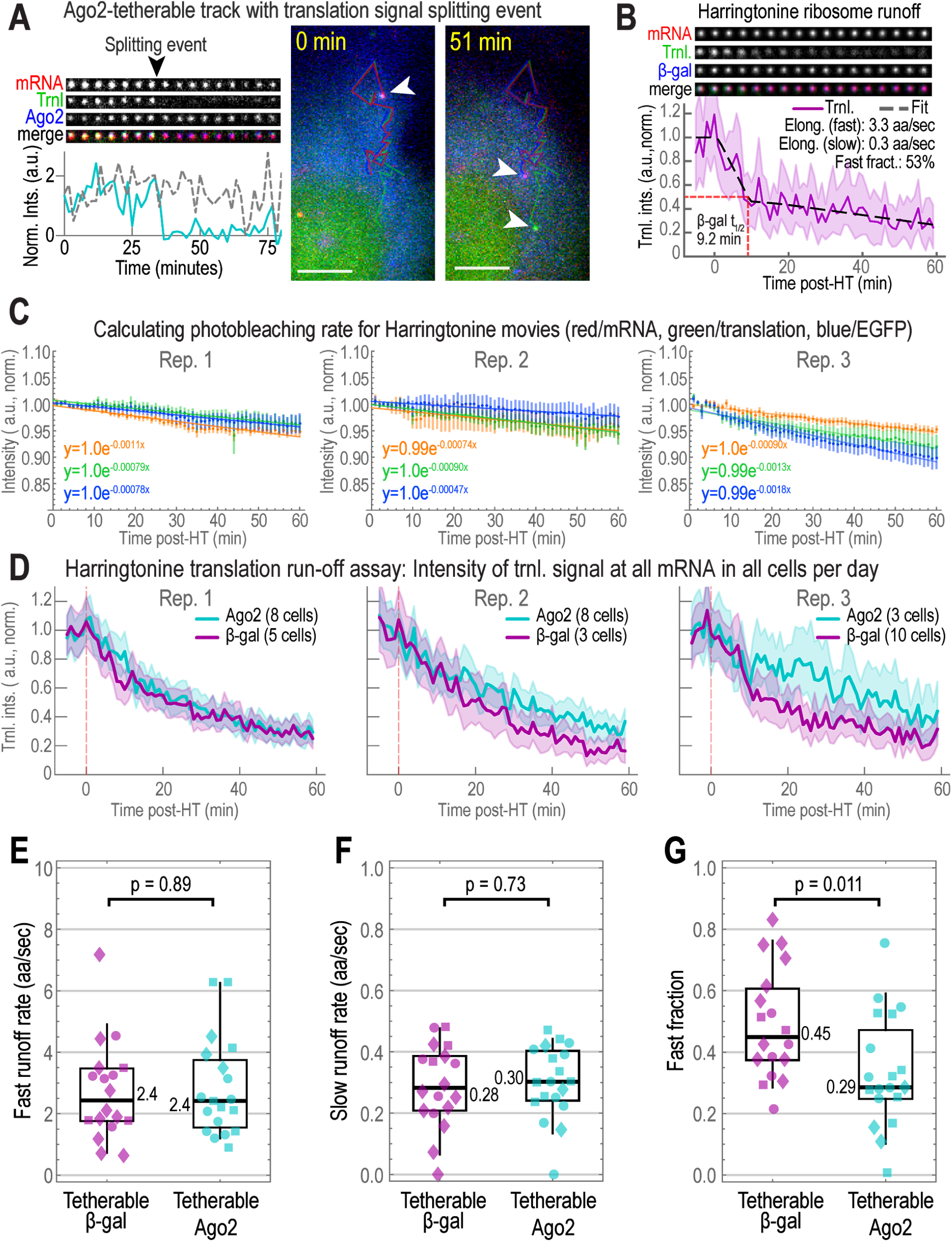
Example event showing the splitting of mRNA and translation signals and Harringtonine experiments analyses. **A** One of two example tracks from **Fig. 4C** where the translation signal split from the mRNA. Plotted as in **Fig. 4D**. Right, two sample images are shown with the tracks overlaid (mRNA, red; translation, green; Ago2, blue). Arrows indicate the current location of the biosensor at the displayed times (yellow text). Scale bar, 5 μm. **B-F** Harringtonine (HT) ribosome runoff assays were performed on 18 cells expressing tetherable Ago2 and 17 cells expressing tetherable β-gal from 3 replicate experiments. HT was added to cells at Time = 0. **B** Average single-molecule signals from all detected TnT biosensors in a single representative cell expressing tetherable β-gal and exposed to HT (Time = 0), plotted as in C. The average translation signal (solid line) showing ribosome runoff was fit to a biphasic model (dashed line; fitted parameters displayed). Red line denotes the runoff halftime t_1/2_. **C** Signal photobleaching over time in HT replicate experiments was measured, plotted, normalized to start at 1 and fit to a single exponential decay (mRNA, red; translation, green; tethering, blue; error bars show SEM; fit denoted with solid lines). For each replicate experiment, mean background-subtracted, whole-cell intensity was measured at each timepoint and plotted. **D** Average signals from TnT biosensors through time for each replicate HT-runoff experiment, plotted as in **Fig. 4C** (Ago2; cyan, β-gal, purple). HT was added at Time = 0, denoted by vertical, dashed red lines. Shaded regions, 95% CIs. **E-G** Box plots showing the fitted parameters from the biphasic HT-runoff model (left, fast elongation rate in amino acids per sec, aa/sec; middle, slow elongation rate; right, fast fraction; shapes denote 1 of 3 replicate experiments). P values were calculated using the Mann-Whitney test.

**Supplemental Figure 5.**
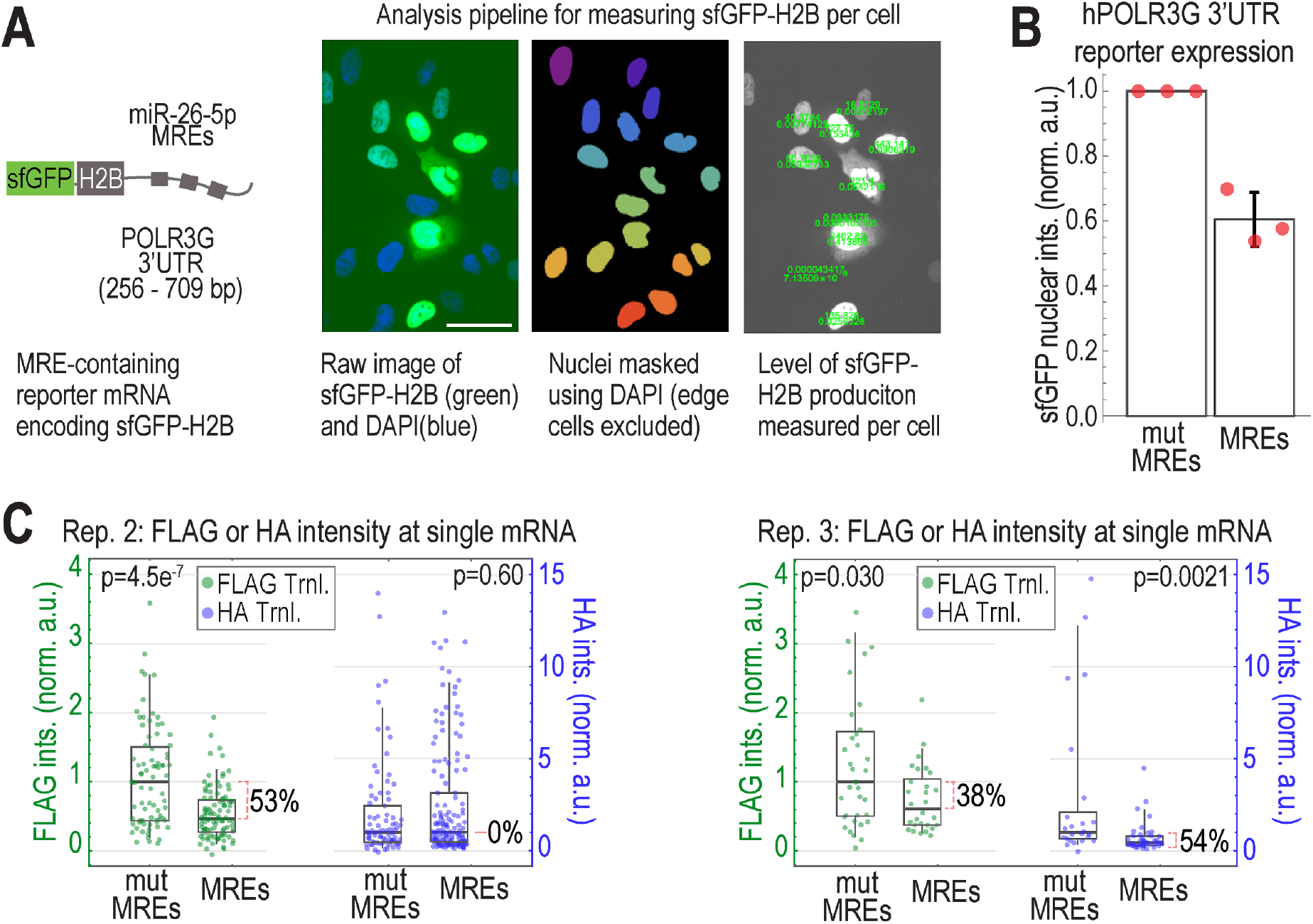
miRNA-directed Ago targeting represses translation at the whole-cell and single-mRNA levels. **A** Schematic of the MRE-containing sfGFP-H2B reporter and analysis pipeline. Scale bar, 25 μm. **B** The nuclear GFP intensity of cells expressing sfGFP-H2B-hPOLR3G 3’UTR with endogenous or mutated (mut) miRNA Response Elements (MREs) was quantified. Points represent the medians per replicate experiment. N = 366, 325, and 1258 nuclei for each replicate experiment for the mutated-MRE-containing reporter (mut MREs) and N = 883, 458, and 1803 nuclei for each replicate experiment for the MRE-containing reporter (MREs). **C** Box plots showing two additional replicate experiments like the one shown in **Fig. 5C**. Left, N = 79 mut and N = 99 MRE green spots for FLAG; N = 85 mut and N = 153 MRE blue spots for HA. Right, N = 35 mut and N = 28 MRE green spots for FLAG; N = 24 mut and N = 44 MRE blue spots for HA.

## Notes

https://doi.org/10.6084/m9.figshare.c.5395800.v2

### Competing Interest Statement

The authors have declared no competing interest.

